# Remote sensing of emperor penguin abundance and breeding success

**DOI:** 10.1101/2023.08.24.554580

**Authors:** Alexander Winterl, Sebastian Richter, Aymeric Houstin, Téo Barracho, Matthieu Boureau, Clément Cornec, Douglas Couet, Robin Cristofari, Claire Eiselt, Ben Fabry, Adélie Krellenstein, Christoph Mark, Astrid Mainka, Delphine Ménard, Jennifer Morinay, Susie Pottier, Elodie Schloesing, Céline Le Bohec, Daniel P. Zitterbart

## Abstract

Emperor penguins (*Aptenodytes forsteri*) are under increasing environmental pressure. Monitoring colony size and trends of this Antarctic seabird relies primarily on satellite imagery recorded near the end of the breeding season, when illumination levels are sufficient to capture images, but colony occupancy is highly variable. To correct population estimates for this variability, we develop a phenological model that accurately predicts the number of breeding pairs and fledging chicks, as well as key phenological events such as arrival, hatching and foraging times, from as few as six data points from a single season. The ability to extrapolate occupancy from sparse data makes the model particularly useful for monitoring remotely sensed animal colonies where ground-based population estimates are very rare or unavailable.

**Teaser:** The Emperor penguin becomes the Southern Ocean’s canary in a coal mine through remote sensing its annual breeding success.

## INTRODUCTION

Emperor penguins (*Aptenodytes forsteri*) breed on land-fast sea ice, making them particularly vulnerable to the impact of global warming (*1–6*). For breeding, stable land-fast sea ice (or fast ice), which is sea ice anchored to land, ice shelves, or grounded icebergs, is required (*5, 7*). The extent of the Antarctic fast ice is predicted to rapidly decline in the coming decades (*8*), and the species is predicted to become extinct by the end of this century. A significant decline of sea ice could already be observed in 2023 (*9*). Despite this, the majority of emperor penguin colonies remain insufficiently studied due to the remoteness and harsh environmental conditions of their habitat (*5, 10*). New strategies to better understand this species and their responses to changing environmental conditions are urgently needed.

Of the 61 currently known emperor penguin breeding colonies (*5*), ground truth population counts conducted during the winter, at regular, frequent intervals, are only available for the colonies at Pointe Géologie in Adélie Land (average population size of 3900 breeding pairs)(*11, 12*) and Atka Bay in Dronning Maud Land (average population size of 8600 breeding pairs)(*13*). Only from these closely-monitored colonies, precise parameters of the species’ reproductive cycle and life cycle have been recorded.

Emperor penguins return to their breeding colony at the onset of the lightless Antarctic winter, between late March and early May, to begin their annual breeding cycle. After a period of courtship, copulation, and egg-laying lasting 6 to 10 weeks, females leave the colony to forage and replenish their body reserves at sea, while the males remain to incubate the single egg for an average of 64 days (*14, 15*). The birds limit their body heat loss during incubation by forming tight groups, so called huddles (*14, 15*). Chicks hatch in winter, i.e. between July and August, and are not thermally independent during the first 6 to 7 weeks of life, therefore one parent always stays with the chick while the other forages. After this brooding period, chicks are left alone at the colony, forming crèches with the other chicks, so that both parents can forage simultaneously to satisfy the chick’s growing demands. Parents feed their chicks between 7 and 12 times until fledging between November and January (*14, 15*), just before the land-fast sea ice begins to break up. The duration of these foraging trips declines from 15-29 days after hatching to less than 10 days before fledging (*16–21*).

Since the majority of colonies are not surveyed by ground-based observations, satellite-based surveys provide the bulk of available datasets for population size estimation. As the resolution of satellite imagery has improved over time, satellite-based surveys hold the greatest potential for estimating global populations and detecting trends (*5, 22, 23*). However, even at the highest resolution currently feasible, which is on the order of 0.5 to 1 m/pixel, satellite-based surveys cannot yet provide measurements at the individual level, but rather estimates of the number of animals based on the area occupied by the colony.

The viability of using colony area to estimate abundance suffers from uncertainties introduced by the imaging process (satellite off-nadir, sun azimuth, and sun elevation (*22*)), the conversion of colony areas to numbers of individuals, and large fluctuations of colony occupancy due to the species’ phenology. The conversion of colony area to numbers of individuals requires knowledge of the average area number density (number of animals per square meter), which is subject to hourly fluctuations, as the animals regulate their body temperature by huddling together or loosening up, depending on weather conditions such as temperature, wind speed, solar radiation, and humidity (*22, 24, 25*). Moreover, the occupancy fluctuations due to the annual phenology pattern is modulated by many factors such as sea ice extent and prey availability, which both influences the duration of foraging trips and the foraging success (*19, 26*).

Due to polar night, satellite images are not available during the incubation stage (June to July), when the least variation in the numbers of individuals is expected. Instead, usable satellite images can only be obtained between September and January, when chicks and only a fraction of the adults are present at the colony. The low resolution of the images does not allow for distinguishing between adults and chicks. For these reasons, satellite-based surveys suffer from large uncertainties when estimating the number of breeding pairs. Moreover, the breeding success of a colony based solely on the number of surviving chicks can not be determined (*22, 23*). Currently, satellite-based surveys may be sufficient to roughly gauge population size and long-term trends, but not to assess short-term selective forces impacting breeding success, unless the colony has completely disappeared (*27*).

The aim of this study is to develop a method to compensate for the uncertainties of satellite-based surveys and to provide an estimate of the annual number of breeding pairs as well as the annual breeding success of a colony, based on the colony area measured during the austral spring and summer (September to December). To achieve this, we combine data from ground-based manual counts and images with a mechanistic phenological model for emperor penguin population counts. The model parametrizes the phenological annual breeding pattern and, when applied to data, infers key phenological and breeding parameters such as arrival time at the colony for breeding, breeding cycle duration, number of breeding pairs, number of hatched chicks, and number of fledgling chicks. Furthermore, we demonstrate that the model remains accurate even when the model parameters are estimated from sparse population counts, and even when these counts are obtained over only a short time window. We then apply our model to satellite-based data to estimate animal numbers and breeding success.

## RESULTS

### Phenological model

We count the number of emperor penguin adults and chicks on a weekly basis at Atka Bay (AB, 70° 40’S, 8° 16’W) over 3 breeding seasons (2018 to 2020), and at Pointe Géologie (PG, 66° 40’S, 140° 01’E) over 10 breeding (2012 to 2021). The data reveal a characteristic pattern of animal counts depending on annual cycles of incubation, guarding, foraging at sea, and feeding chicks at the colony (Fig. 1).

**Fig. 1.**
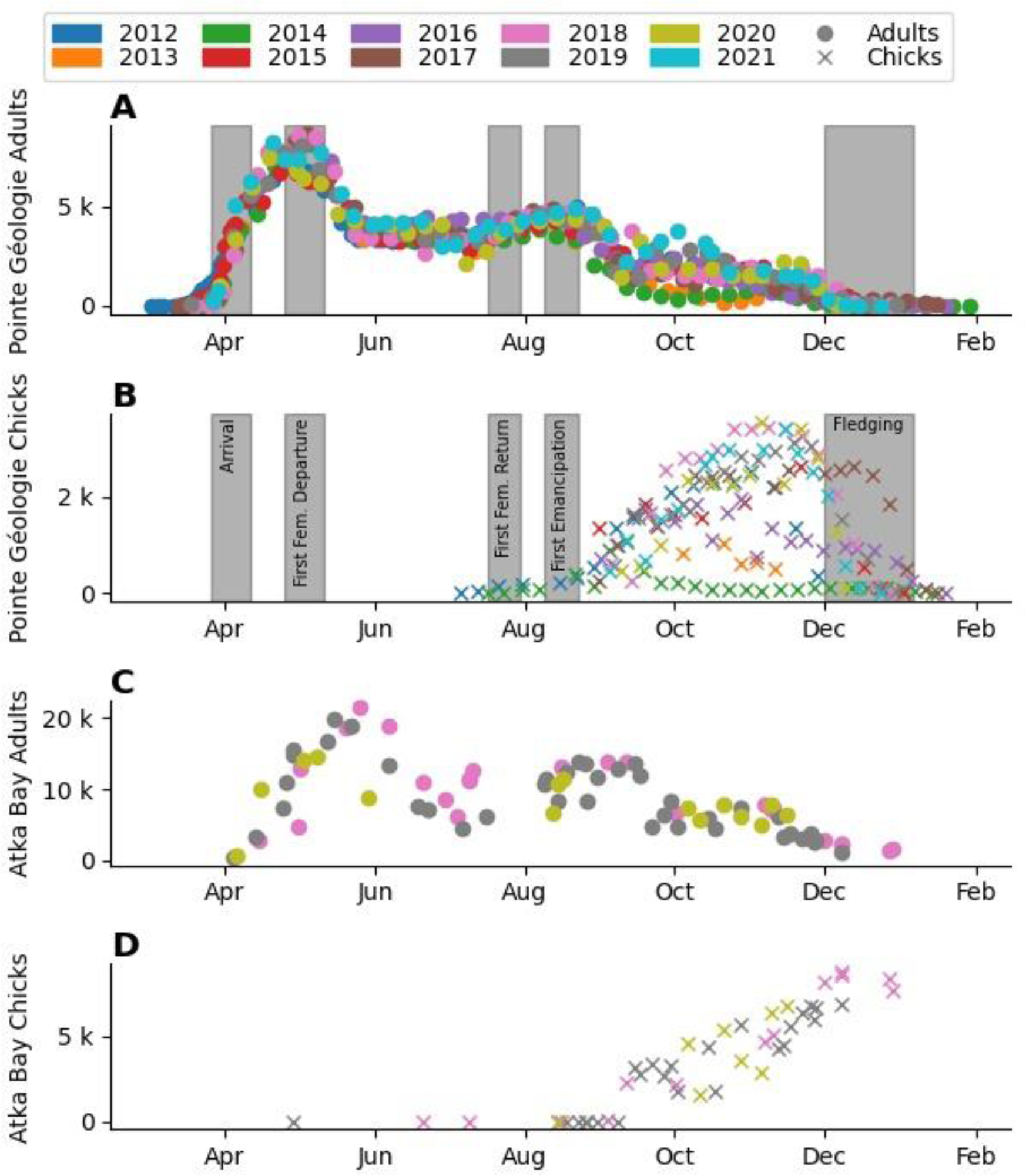
Manual counts of the number of individuals. Number of individuals at Pointe Géologie (**A**,**B**) and Atka Bay (**C**,**D**) colonies. Circles indicate counts of individual adults (**A**,**C**), crosses indicate counts of individual chicks (**B**,**D**). Colors denote different breeding seasons. Gray bars show the time range of key phenological events extracted from manual observations at Pointe Géologie (**A**,**B**). Note that the number of chicks increases between August and November as the chicks become thermally independent and therefore less occluded.

Based on known patterns of the species’ breeding activity (*16–21*), we develop a mechanistic phenological model that describes the observed colony abundance during a typical breeding season (from March 1 to February 28 of the next year). This model describes the presence/absence pattern for a breeding pair and its chick based on the following set of parameters: time of arrival at the colony site, courtship duration, first female absence duration after laying, foraging trip duration, and period at the colony to feed the chick. The model then calculates the total number of females, males, and chicks within the colony at any given time point from the presence/absence patterns. We fit the model parameters to the observed number of adults, using a Markov chain Monte Carlo approach, which iteratively optimizes the parameters and returns a distribution for each model parameter as well as a distribution for the number of chicks and adults for each day of the breeding season.

To validate the model, we compare the observed numbers of individuals to the model predictions (see Fig. 2). Because the count data usually vary on a logarithmic scale, we report not arithmetic, but geometric errors: for example, a ±25% geometric error corresponds to an overestimate of 25% (multiply by 1.25) or an underestimate of 20% (divide by 1.25). We find an average geometric error of 16% for all data points (AB: 23%, PG: 15%). 81% of the observations fall within one standard deviation around the model estimate (AB: 83%, PG 81%). The coefficient of determination (R^2^ value) between model and data is 0.91 for all data points (AB: 0.73, PG: 0.95).

**Fig. 2.**
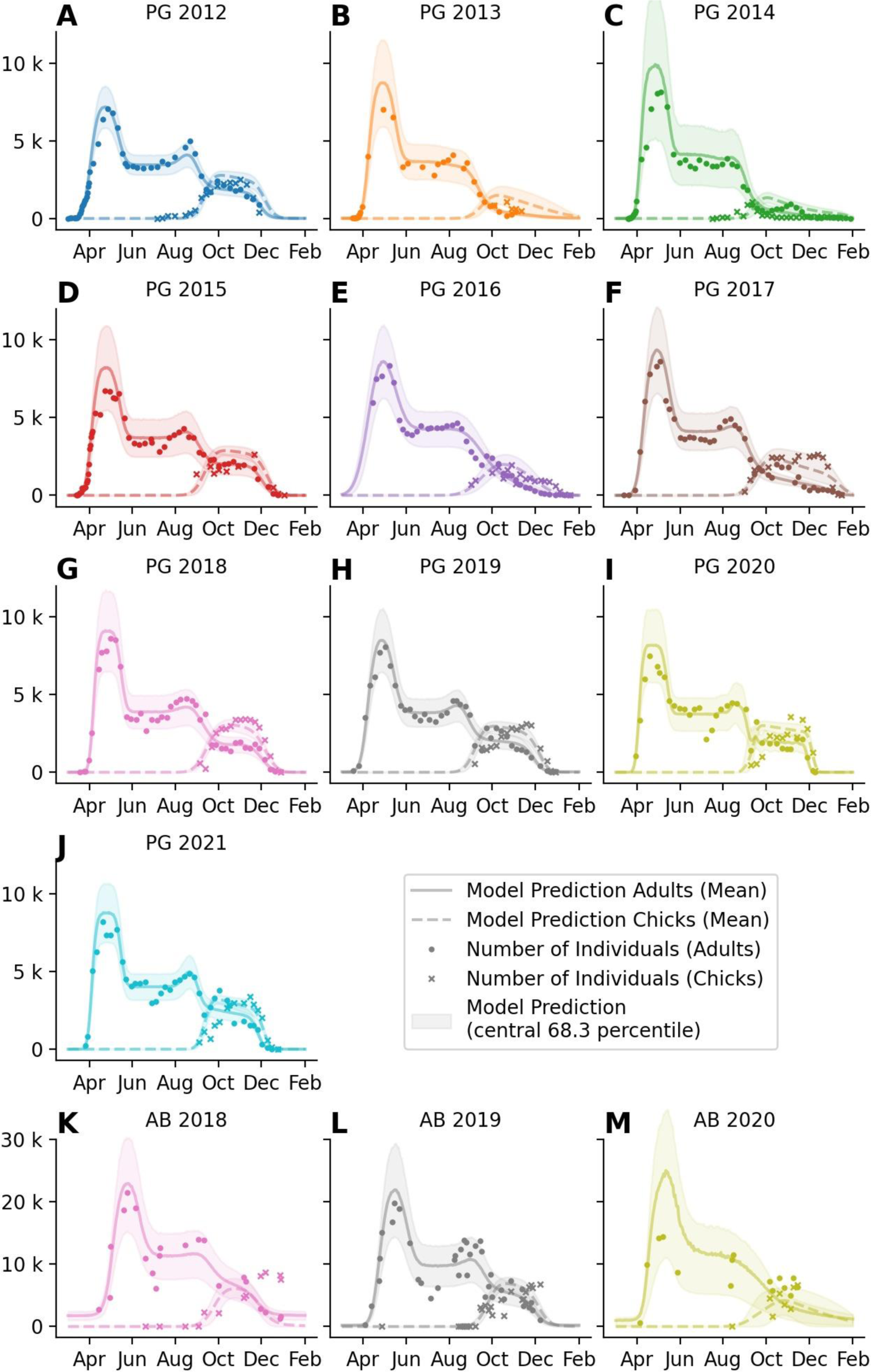
Phenological model fit to manual counts. Overlay of data and model output for all observed seasons at Pointe Géologie (10, **A-J**) and Atka Bay (3, **K-M**) colonies. Individual panel titles indicate colony (PG = Pointe Géologie, AB = Atka Bay) and season. Y-axes show the number of individual adults (circles) and chicks (crosses). Solid and dashed lines show the model prediction for the adult and chick counts, respectively. The shaded areas indicate the ±1-sigma confidence interval of the model prediction. Colors indicate the season as in Fig. 1, 3 and 8.

The model also provides date estimates for phenological events such as arrival, first departure and first return of the females after laying, chick emancipation (when the chick is thermally independent to join the other chicks in crèches), and fledging (when the chick has molted and leaves the colony for the first time). We compare these estimates with ground-based observations for Pointe Géologie (Fig. 3C, Table S11). The timing of events as estimated by the model shows good agreement with the observed timing of events (R^2^=0.98, average absolute error of 10 days) when pooled over all years. For individual years, we find good agreement of the model prediction for all events. Note that a difference of 14 days arises between model prediction and ground truth times for female first departure and chick emancipation, because these events are recorded in the field as the time of the first observation of the respective behavior, whereas the model predicts the central moment of the event.

**Fig. 3.**
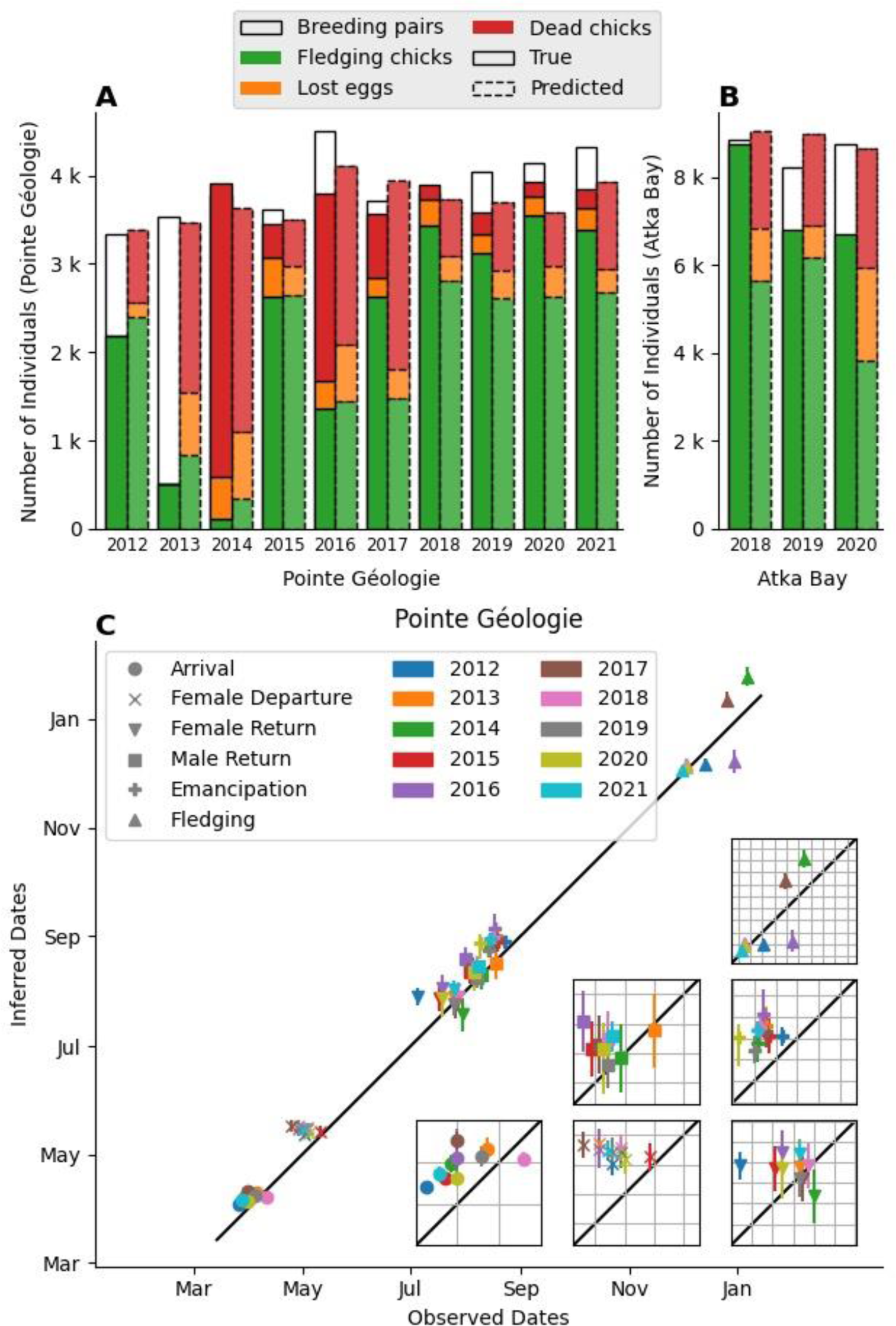
Model estimates of key phenological parameters. (**A-B**) Observed and predicted breeding success by year and colony. The solid black and dashed black bars show the true and predicted total number of breeding pairs. The colored areas split the bars into fledged chicks (green), dead chicks (red), and lost eggs (orange). The predicted numbers add up by model definition while the ground truth values do not, due to counting inaccuracy. The solid line separates between Pointe Géologie and Atka Bay data, and the left and right Y-axis. (**C**) Time of predicted phenological events vs observed phenological events. Each point corresponds to one season, the y-bars show the 1-sigma confidence interval. Colors indicate the seasons. The black line shows the line of identity. Insets show the same data, zoomed in for better visibility, to show inter-annual variation. Gridlines indicate weeks. Note that systematic shifts between observed and estimated dates arise from the difference of first occurrence (manual observations) and central event date (model estimates).

### Predictive capacity of the phenological model

A key feature of our model is its ability to predict the annual breeding success expressed as the number of fledging chicks relative to the annual number of breeding pairs. Because we only fit the model parameters based on the number of adults, we can assess the predictive power of the model by comparing the model estimate of the number of chicks with the actual number of chicks observed. We find an overall R² value of 0.45, with R²=0.64 for Pointe Géologie and R²=0.12 for Atka Bay (Fig. 2). The average geometric error is 41% for all data, with 51% for AB data and 39% for PG data. The large relative error for Atka Bay is attributable to the unusually (compared to other AB and PG seasons) high number of chicks that were still present late in the 2018 season (see Fig. 2K), which the model fails to predict. Further, the average number of sample counts per season for Atka Bay (26 counts) is lower than for Pointe Géologie (44 counts). In addition to the number of alive adults and chicks per day, the total number of fledging chicks, dead chicks and lost eggs are available from Pointe Géologie but not from Atka Bay. For Pointe Géologie, the model predicts the number of fledging chicks and dead chicks with R² values of 0.74 and 0.32, and average geometric errors of 25% and 56%, respectively (Fig. 3A,B). The model predicts the number of dead eggs with a mean absolute error of +/- 190 animals.

### Phenological parameters

The model describes each season and each colony separately. Therefore, we extract 13 sets (from 13 breeding seasons) of 14 parameters (*BP, H, F, NB, t_0_, Δt_0,_ m, b, Δb, c_max_, c_min_, s_max_, s_fem_, s_min_*) that provide insight into the interannual and inter-colony variation of the phenology (Fig. S2 in the Supplement).

Most prominently, the arrival time (*t_0_*) at Pointe Géologie (April 7 +/- 3 d) is significantly (P<0.01) earlier than at Atka Bay (April 27 +/- 7 d). The number of breeding pairs (*BP*) also shows significant (P<0.01) differences between those two colonies (AB: 8600 breeding pairs on average, PG: 3900 breeding pairs on average). In addition, we find statistically significant (see S8 in the supplement) interannual variance for the number of breeding pairs (*BP*, AB: +/- 620), fledging success (*F*, PG: +/-0.66), time of arrival (t_0_, AB: +/-12.0 d, PG: +/-4.6 d), width of arrival date distribution (*Δt_0_*, PG: +/-3.8 d), width of female return date distribution (*Δb*, PG: +/-7.0 d), minimum time at sea (*s_min_*, PG: +/-6.4 d), and maximum time at sea (*s_max_*, +/- 6.7 d). Note, however, that the standard deviation for the duration of each event is smaller than the sampling interval of 7 days for the ground truth data. Consequently, the biological significance of these interannual variations cannot be reliably verified.

We further investigate the correlation between model parameters versus breeding success, as estimated by the ground truth ratio of fledging chicks to breeding pairs. We find significant (P<0.02, n=13 seasons) correlations between breeding success and the foraging trip duration during the crèching period with the following R² values: 0.54 (*s_max_*), 0.52 (*s_min_*), 0.46 (*c_min_*) (see Table 1, Fig. S5 in the Supplement). We then sum the duration of all foraging/feeding trips to obtain the total time spent at sea or at the colony during the crèching phase (Fig. 4). We find a significant (P<0.03) correlation between breeding success and total time spent at sea (R²=0.55) or time spent at the colony (R²=0.39). Noteworthy, a longer total time spent at the colony correlates with a higher breeding success, while a longer total time spent at sea correlates with a lower breeding success.

**Fig. 4.**
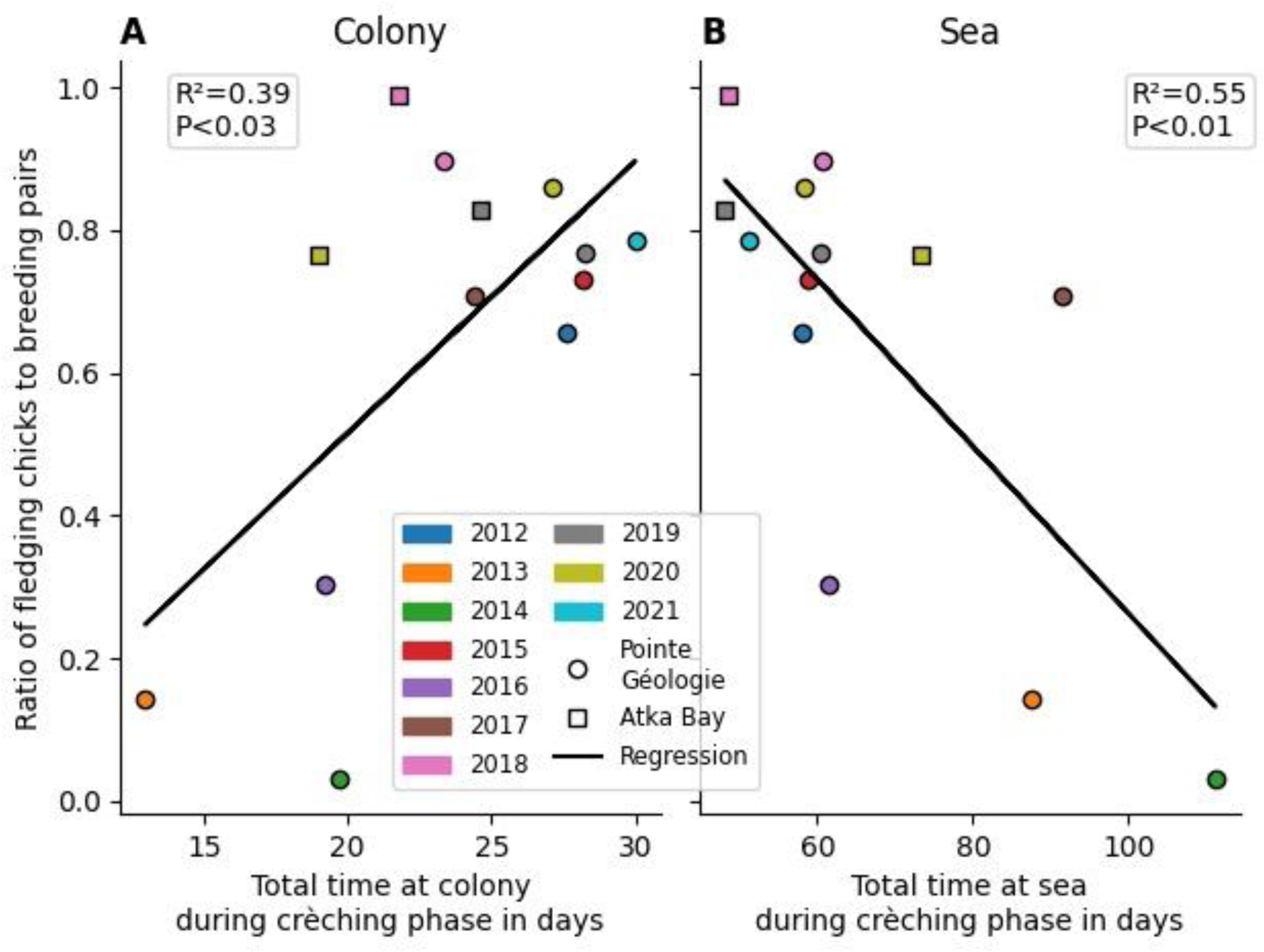
Correlation between foraging pattern and breeding success. The x-axis shows the total number of days spent at the colony (**A**) or at sea (**B**) during the crèching phase as predicted by the model, averaged for females and males. The y-axis shows the ratio of fledging chicks to breeding pairs (breeding success) from manual observations. Every dot (Pointe Géologie) and square (Atka Bay) denotes one season. The black lines show the regression lines for each point cloud (Pointe Géologie and Atka Bay merged). We find a significant (n=13, P<0.03) positive correlation between breeding success and time at colony, and a significant (n=13, P<0.01) negative correlation between breeding success and time at sea.

**Table 1.** Phenological Model Parameters. Parameter names, descriptions, and numerical ranges.

| Symb<br>ol | Name | Lower<br>Limit | Upper<br>Limit | Unit | Fit per<br>year<br>/<br>global | Description |
| --- | --- | --- | --- | --- | --- | --- |
| $t_0$ | arrival | 50 | 150 | days | year | Peak of animal arrivals at the |

| Symbol | Name | Lower Limit | Upper Limit | Unit | Fit per year / global | Description |
| --- | --- | --- | --- | --- | --- | --- |
|  | date | (Feb 1) | (May 30) |  |  | colony |
| $\Delta t_0$ | arrival date standard deviation | 4 | 14 | days | global | Standard deviation of arrival dates |
| $m$ | courtship duration | 28 | 42 | days | global | Average duration between arrival and first departure of females |
| $b$ | female absence duration | 50 | 100 | days | global | Average duration between first departure and return of females |
| $\Delta b$ | female absence duration standard deviation | 0 | 14 | days | global | Standard deviation of duration of female absence |
| $BP$ | number of pairs | 2.000 | 15.000 | pairs | year | Number of pairs that mate and produce an egg. |
| $NB$ | non breeder to breeding pair ratio | 0 | 1 | 1 | year | Ratio of birds that do not find a breeding partner and leave the colony before the breeding period, to the total number of breeding pairs |
| $H$ | hatching success ratio | 0 | 1 | 1 | year | Ratio of pairs that repeatedly return after the breeding period, relative to the number of breeding pairs at the beginning of the breeding season |
| $F$ | fledging success ratio | 0 | 1 | 1 | year | Ratio of pairs that repeatedly return until the end of the chick rearing phase, relative to the number of breeding pairs at the beginning of the breeding season |

| Symb<br>ol | Name | Lower<br>Limit | Upper<br>Limit | Unit | Fit per<br>year /<br>global | Description |
| --- | --- | --- | --- | --- | --- | --- |
| $C_{max}$ | maximu<br>m time<br>at<br>colony<br>per trip | 1d | 5d | days | year | Time one parent spends at the colony per foraging trip during the guarding phase and at their first foraging trip. |
| $C_{min}$ | minimu<br>m time<br>at<br>colony<br>per trip | 1d | $C_{max}$ | days | year | Time one parent spends at the colony on their last foraging trip. |
| $S_{max}$ | maximu<br>m time<br>at sea<br>per trip | $C_{max}$ | 21d | days | year | Time one parent spends away from the colony per foraging trip during guarding phase and at their first trip after chick emancipation. This includes foraging and commuting time |
| $S_{min}$ | minimu<br>m time<br>at sea<br>per trip | $C_{max}$ | $S_{max}$ | days | year | Time one parent spends away from the colony per trip on their last foraging trip. This includes foraging and commuting time |
| $S_{fem}$ | time at<br>sea at<br>first trip<br>(only<br>females) | $C_{max}$ | $S_{max}$ | days | year | Duration of the first foraging trip of females after hatching. It is shorter or equal than the following trips. |

### Density estimation from ground-based imagery

To estimate penguin abundance and breeding success based on ground-based imagery, we need to multiply the measured colony area (ground area covered by penguins) with an estimate of the colony density (number of animals per area) to obtain the number of individuals. To estimate the colony density, we use a so-called windchill model that predicts colony density from locally measured meteorological variables, specifically air temperature, wind speed, solar radiation, and relative humidity, as described in (*25*). All of these meteorological variables affect the animals’ heat balance and contribute to an apparent or perceived temperature, analogous to the windchill-corrected temperature reported in US-American weather forecasts. If the perceived temperature falls below a critical temperature, the animals tend to form huddles. Accordingly, the model requires a critical temperature as a further model parameter. Specifically, the critical temperature corresponds to the apparent temperature at which the average density of the animals within a colony reaches half of the maximum packing density in a dense huddle of 12.8 animals per m^2^.

To obtain the windchill model parameters, we manually select the colony covered area from ground-based images of Pointe Géologie and Atka Bay (*13, 28*) using the image annotation software Clickpoints (*29*), project the resulting polygons to the top view projection (*30*), and calculate the total occupied area in m² (Fig. 5A & 5B). From this, we compile a dataset of 538 measurements of colony area, number of individuals (chicks and adults), and meteorological variables (temperature, wind speed, solar radiation, and humidity) acquired at each colony between September and December of three (AB) and four (PG) seasons. From the correlation of colony density fluctuations (colony area divided by the number of individuals) with fluctuations in meteorological variables, we obtain the windchill model parameters (Fig. 5C-F, Fig. S17).

**Fig. 5.**
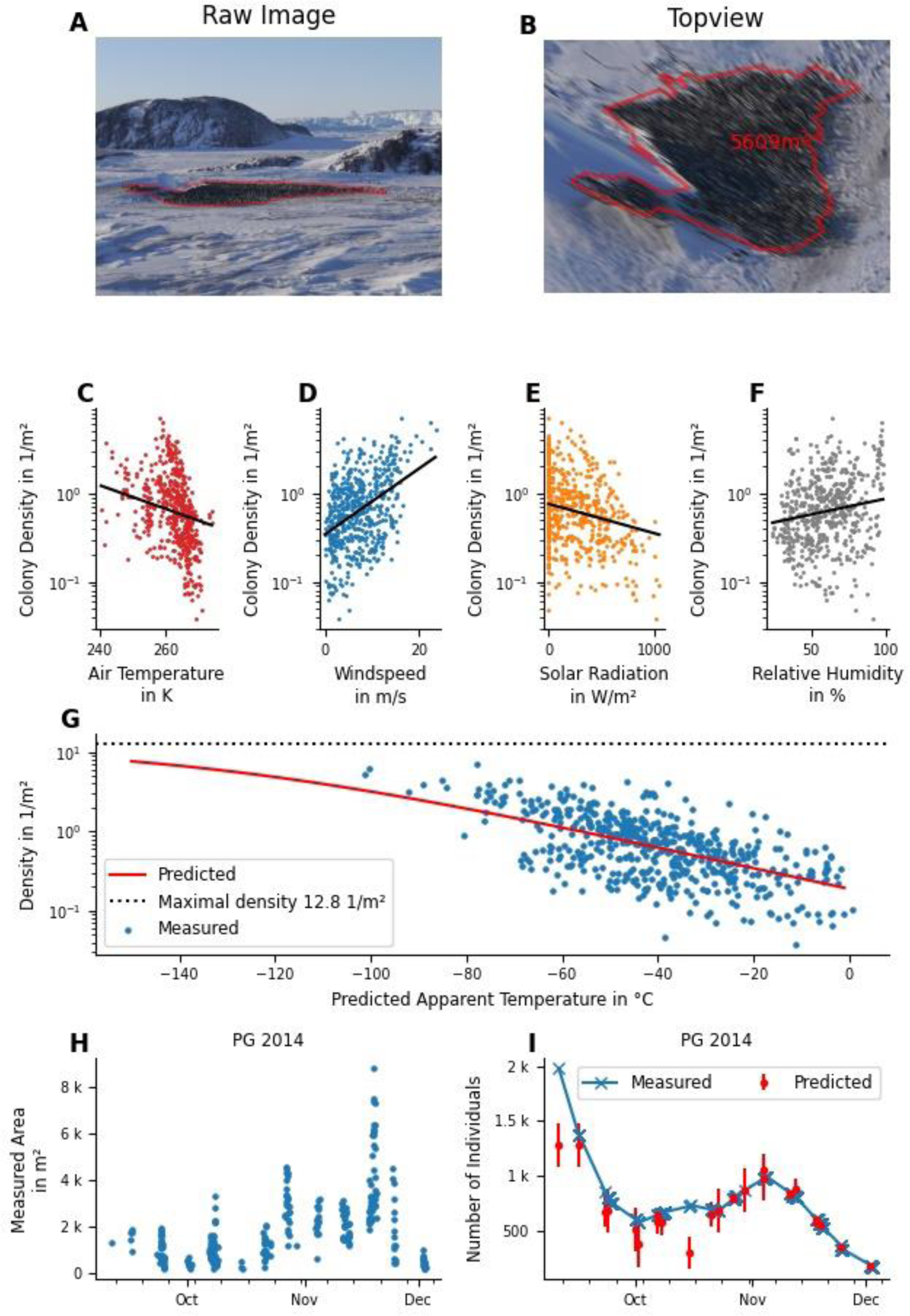
Description of the windchill model. (**A**) Example of an image recorded by the automatic time lapse camera on 2017-08-22 04:00:00 UTC at Pointe Géologie. The manually-annotated outline of the colony is highlighted with a red polygon. (**B**) Projected top view of the image shown in (A). The number indicates the area covered by the colony. (**C-F**) Correlation of measured colony density with the meteorological values: air temperature (C), windspeed (D), solar radiation (E), and humidity (F). The y-axis shows the colony density in penguins per square meter, the x-axes show the respective meteorological variables. Each dot represents one image. The black lines show the corresponding log-linear regression line. The slopes correspond to the model parameters. (**G**) Dependence of measured colony density on apparent temperature T_A_. Each dot represents the data from one image and time point. The red line shows the model prediction (fit of the sigmoidal function (Eq. 2) to the data). (**H**) Surface area covered by the colony (in square meters) for Pointe Géologie between September 1 and December 31 in 2014 over time, estimated from ground-based images. The observed short-term variance (seen as vertical stacks in the data points) are due to daily variation in colony area, driven by environmental parameters. (**I**) Measured (blue crosses) and predicted (red dots) animal count over time. Predictions are based on the measured areas shown in (**H**) for Pointe Géologie between September 1 and December 31 in 2014 multiplied with the density as predicted by the windchill model.

The windchill model predicts the measured colony density with an R² value of 0.32 and an average geometric error of 39%. This error may seem large, but when we multiply the density predicted from the model with the measured area in order to obtain a predicted number of individuals, this accurately matches the actual animal counts with an R² value of 0.93 and an average geometric error of 11%. Moreover, we find that 60% of the individual count data fall within the predicted 1-sigma interval (Fig. 6F). Fig. 5H and 5I show an example of the measured colony area and predicted number of individuals at Pointe Géologie in 2014.

**Fig. 6.**
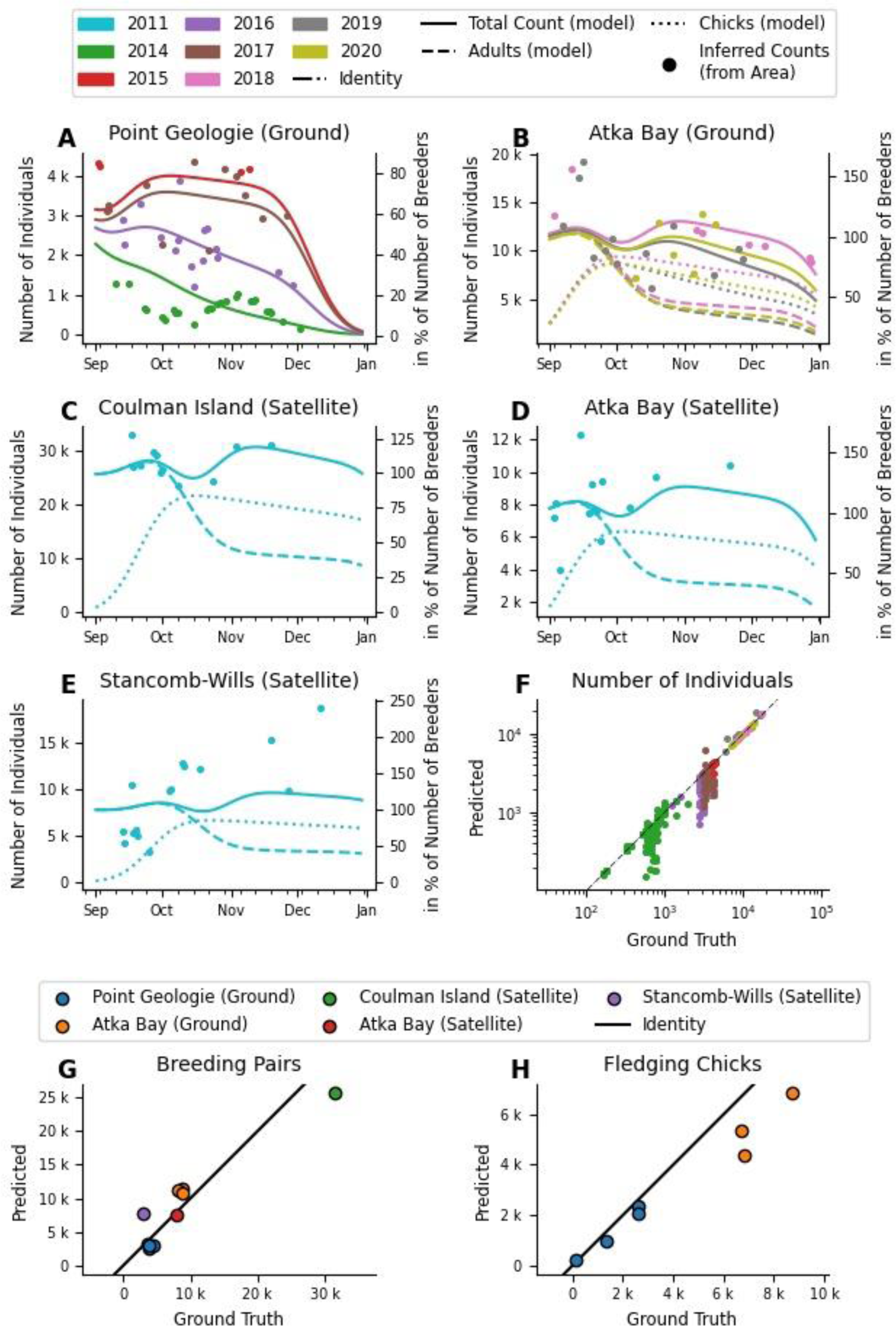
Application to satellite data. (**A-E**) Fit of the phenological model to individual counts inferred from measured colony area and the windchill model predictions (dots). Lines show the phenological model predictions for adults (dashed), chicks (dotted), and total counts (solid). In (A), adult and chick counts are omitted for readability. Those data are provided in Supplement S7. Colors denote seasons. (**F**) Comparison of ground truth counts and counts inferred from colony area and windchill model predictions for Pointe Géologie and Atka Bay over 7 seasons. Colors denote seasons. Note: Pointe Géologie data covers 2012-2017, Atka Bay 2018-2020, therefore colors are also colony specific. The back line shows identity. (**G**,**H)** Comparison of ground truth and phenological model predictions for the number of breeding pairs (G) and fledging chicks (H). Ground truth values for Pointe Géologie and Atka Bay (Ground) are from manual counts, all others are from (*23*). Note that for satellite data, there is no ground truth for the number of fledging chicks.

The windchill model allows us to correct for weather-induced fluctuations in colony density, which is important when computing the number of animals from colony area. Fig. 5H shows fluctuations in colony area for Pointe Géologie between September 1 and December 31. Accordingly, we find large fluctuations of colony density over the course of a day (e.g. from 0.20 to 2.41 animals per m² on 2014-10-08, with a total range between 0.03 and 7.00 animals per m² during the whole observation period; Fig. 5G). These large density fluctuations explain the uncertainties of satellite-based counts when a constant density of 0.93 animals per m^2^ (*23*) is assumed.

### Application to satellite imagery

To showcase an application, we use previously published data of colony area from the population of Coulman Island (CI), Atka Bay (AB), and Stancomb-Wills Glacier (SW) in 2011 (*22*). We obtain the local meteorological data at the time of the satellite image acquisition from (*31*) and, based on the windchill model, predict a colony density between 0.6 animals/m² to 1.8 animals/m². After multiplying the colony area with the colony density, we obtain animal counts, which are the sum of chicks and adults present at the colony. Finally, we fit the phenological model to the counts to obtain the number of breeding pairs and the number of fledging chicks for each year. The results are summarized in Table S13. For these three colonies, low quality ground truth data for the number of breeding pairs exist, albeit not from 2011 but from earlier years (*23*). Nonetheless we find an average geometric error of 28% and an R² score of 0.88.

We also benchmark this approach using ground-based images from AB and PG. The procedure is analogous: we estimate the colony area from the images, use the windchill model to estimate colony density based on meteorological data from station observatories, multiply both numbers to estimate the number of animals, fit the phenological model to the counts, and arrive at an estimate for the number of breeding pairs and fledging chicks for each year that we can compare with ground truth counts. We find good agreement, with an average geometric error of 21%, an R² value of 0.92, and 74% of the ground truth data being within one standard deviation around the model estimate (Fig. 6A-F).

## DISCUSSION

### Phenological model performance

We present a mechanistic model that estimates phenological information from incomplete time series of animal counts in emperor penguin colonies. The phenological information predicted by the model include time of first arrival at the colony site to breed, number of breeding pairs, number of hatched chicks, number of fledged chicks, timing of foraging trips, duration of courtship, and female absence (during incubation). Our phenological model is able to recapitulate the temporal fluctuations of the weekly individual counts at two colonies (Atka Bay and Pointe Géologie) over several seasons, and it accurately predicts the number of chicks per day and the number of fledging chicks solely based on the counts of adult animals.

The model accurately predicts the number of breeding pairs and number of fledging chicks for both colonies. Predicted and true counts of chicks and adults per sampling day align less well at Atka Bay than at Pointe Géologie, probably due to a lower number of sample counts for Atka Bay than for Pointe Géologie (26 vs 44 counts per season). Our model is therefore robust in detecting the main features of the phenological pattern, while struggling with very fine predictions like weekly counts, when the underlying data are sparse. We attribute this to the probabilistic inference method that blurs the parameter space when confronted with noisy and sparse data.

The model has difficulty in years with highly unusual presence/absence patterns at specific times in the breeding cycle. For example, in 2017 at Pointe Géologie, the number of adults present during the chick rearing period was much lower than the number of chicks. The model (which is informed only about the number of adults but not the number of chicks) therefore estimated a survival rate of less than 50%, while in reality, significantly more chicks survived (71%). We hypothesize that, in 2017, adults spent extremely long periods outside the colony, reflecting an extended sea ice cover at the end of chick rearing (*32*), while food resources were likely available and abundant near the sea ice edge, enabling them to feed their chick sufficiently until they fledge. Moreover, the model is based on a very limited knowledge and data on the individual trip durations during the chick-rearing phase (i.e. the study of Kirkwood & Robertson (1997), which was carried out over a single year on the Auster and Taylor Glacier and Auster colonies). To refine the model, it would be important to collect foraging information on this critical period of the breeding cycle, over several years and on several colonies facing contrasting environmental conditions.

Overall, the model performs well at predicting the number of fledging chicks and breeding pairs, although it gives large numbers of dead chicks and lost eggs compared to manual counts (Fig. 3A). A part of the discrepancy arises because lost eggs and dead chicks are counted only if they can be found, but are often covered by snow. Therefore, the model may in fact provide more reliable estimates for the number of dead chicks and lost eggs in years with typical phenology.

### Timing in phenological parameters

The model performs well for predicting phenological events on average. We could not quantify how well the model can predict annual variations in phenological events at the same colony, because most events vary less than the resolution of our observations (7 days, see Fig. 3C). Nonetheless, some interesting model predictions emerge. For example, the duration of courtship (*m*) and the females’ first absence from the colony after laying (*b*) (the incubation period for the males), do not show significant variability between years and colonies, which is to be expected because such processes have likely evolved to be tightly regulated at a physiological level specific to the species. We also found a negative correlation of the total time spent at sea with breeding success: long (>10 days) foraging trips at the end of the chick-rearing phase (between September to December) lead to lower (<50%) breeding success, likely because the frequency of feeding events decreases, exposing the chick to a higher risk of starvation in between feeding events. We suggest that increased foraging trip durations are linked to longer distances to the land-fast ice edge, which have been shown to negatively impact fledging success (*32*).

The model-predicted as well as measured arrival times (*t_0_*) and first hatchings (*t_H_*=*t_0_*+*m*+*b*) between Pointe Géologie and Atka Bay differ by approximately 30 days. At Pointe Géologie, the more northerly colony (66.39°S), first hatchings are observed in the first week of July, while at Atka Bay (70.60°S), hatchings are estimated (by our model) to occur in the first two weeks of August. In both colonies, the timing of hatchings correlates with the first sunrise after mid winter (June 29 at Pointe Géologie and July 28 at Atka Bay, Fig. S18). A positive correlation between arrival times to breed and latitudes has been previously reported (*33*). A likely explanation of this correlation is that penguins synchronize the chick rearing phase and therefore their whole annual cycle with the increasing abundance of prey. Prey abundance, in turn, is triggered by a rise in primary production due to an increase in solar radiation and/or a decrease in sea ice cover near the Antarctic continent. A similar synchronization between prey abundance and breeding cycle has been observed in numerous species, including other seabirds (*34*), and is in line with the match/mismatch hypothesis (*35–37*) that states that the reproductive success of a species depends on its ability to match its phenology with that of its prey species.

### Windchill model

We find a relationship between colony density measured from ground-based images and meteorological conditions. This relationship is described by a previously reported model (which for simplicity we refer to as the windchill model) (*25*). We re-trained the windchill model with data from Atka Bay and Pointe Géologie obtained between September and December, as the previously reported model parameters were trained with data obtained during the high-density (up to 15 animals per m²) courtship and incubation phase of the austral winter, in April and May.

The re-trained model can only partially explain the fluctuations in colony density with a correlation of R²=0.32 and 60% of data within the predicted 1 sigma interval, which is slightly higher than the percentage of the original model (50%, (*13*)). This is to be expected because of the mild weather conditions of the austral spring, and because a mixed colony of adults and chicks rarely adopts a configuration where the majority of the animals are found in dense huddles.

### Estimating abundance and breeding success from remote sensing

Satellite images can only be taken during periods when occupancy is most erratic and therefore the most challenging for inferring population size and success. One of the main objectives of our phenological model is to provide more reliable estimates of the number of breeding pairs within a colony from those images. Currently, it is common practice to multiply the colony area *A* (in units of m^2^) as measured from satellite images with a conversion factor *CF* of 0.93 breeding pairs per m^2^ (*23*). However, depending on the weather conditions and the time of year, the colony area and animal density fluctuate, potentially contributing to large uncertainties (*22, 23*). To reduce these uncertainties, we suggest the following approach for a reliable estimate of the number of breeding pairs: (1) Collect several (5 to 10) satellite images from multiple time points *t* (over a time period of 2 months, from October to December) and measure the colony area *A*; (2) Acquire local weather conditions at the time of satellite image capture and estimate colony density ϱ using the windchill model; (3) Compute the total number *N* of animals in the colony (N = ϱ A); (4) Fit the phenological model to *N*(*t*) and extract the number of breeding pairs *BP* and fledged chicks *F*.

To benchmark this method, we estimate the number of breeding pairs from ground-based images of the colony (instead of satellite images) for our two colonies (AB and PG) over a time period of 3 and 4 years, and find an excellent correlation (R²=0.93). Note that if multiple images are recorded on a single day, the predicted counts are averaged in order to reduce sampling bias. Furthermore, we compute a conversion factor *CF* from the number of breeding pairs and the colony area as *CF = BP/A* (see S15 in the Supplement). We obtain a value of *CF =* 1.09 breeding pairs per m², which is close to the conversion factor of 0.93 breeding pairs per m² as reported in (*23*). Note that when we compute the average *CF*, we exclude an outlier from the 2014 season at Pointe Géologie with an unusually low (<5%) breeding success, a very low number of animals in the colony (approximately 500 instead of the usual 1800 animals on average), and a very high density of 5.98 breeding pairs per m². However, this outlier is not excluded when we compute the correlation of the model estimates with manual counts. Taken together, we are convinced that previous estimates of the total number of breeding pairs from satellite images are accurate on average, but individual counts from any given image can be subject to large errors, as was already pointed out in (*22, 23*). The number of surviving chicks that we estimate with our model following the approach outlined above also agrees well with manual counts (R²=0.81 and average geometric error of 26% based on data from two colonies over 3 or 4 years).

We find a rather striking difference in the number of breeding pairs at Atka Bay between the periods 2008-2011 and 2018-2020. In 2008-2011, the estimates range from 7300 (satellite images, Table S13) to 9657 (satellite images, (*23*)), while in 2018-2020, the estimates are around 10,000 (both from ground truth counts and ground-based images). This increase by more than 2,000 breeding pairs is likely due to an immigration of individuals from the neighboring Halley colony that experienced three consecutive years of breeding failure (from 2014 to 2016) and had completely abandoned the colony site in 2016 (*23*).

While the number of breeding pairs provides valuable information about the current status and health of the species, the number of surviving chicks during a breeding period is a more sensitive predictor of future population trends, e.g. due to global change or changes in food supply. As of now, we have not detected a declining trend in the number of surviving chicks at AB or PG over the 2012-2021 period. However, given the recent development of a dramatically decreased sea ice extent around Antarctica (*9*), chances for the long term survival of the species stay dire.

The next milestone will be to apply our method on a long-term circum-Antarctic scale, enabling us to use the emperor penguin breeding success as an early warning indicator. This indicator will alert the scientific community on an annual level about potential breeding failures, allowing us to better correlate such breeding failures with local oceanographic events. Ultimately we aim to provide stakeholders and governments with information on the health of populations and their ecosystems to rapidly implement conservation measures.

## MATERIALS AND METHODS

### Individual counts

The individual counts were conducted at the Pointe Géologie (PG) emperor penguin colony in vicinity to the Dumont d’Urville French research station (66° 40’S, 140° 01’E) and at the Atka Bay (AB) colony in the vicinity to the Neumayer III German research station (70° 40’S, 8° 16’W). The data from Pointe Géologie were collected over 10 years (2012 to 2021), Atka Bay data were collected over 3 years (2018 to 2020) (Fig. 1). Weekly counts were acquired by imaging the colonies with either hand-held (Pointe Géologie) or remote-controlled cameras (Atka Bay, (*13*)) from an elevated position, resulting in one or several panoramic images per day that have sufficient resolution to count individual penguins. For manual counting, we used Adobe Photoshop or Clickpoints (*29*).

Manual observations of phenological events such as time of arrival, female departure, or the beginning of fledging were recorded only at Pointe Géologie. However, some events (e.g. beginning of fledging, see Table S11 in the supplement) could not be recorded in some years for various reasons such as inaccessibility of the colony, occlusion of the colony by geographic landmarks, or a low number of breeding pairs.

The ground truth numbers of breedings pairs at AB and PG were taken from individual counts during the incubation period when only the incubating males are at the colony. Lost eggs/chicks were counted daily at PG. The number of fledged chicks was not counted.

Therefore, we used the last known number of chicks before the onset of fledging as an estimate of the number of surviving chicks.

### Colony area and colony density

Colony area was measured from images after perspective correction. The Pointe Géologie colony was monitored between 2014 and 2017 using automatic time lapse cameras (*28*). For Atka Bay, we used the panoramic images from the seasons 2018, 2019, and 2020. We manually marked the colony boundaries on 538 images (509 of Pointe Géologie, 29 of Atka Bay) acquired between 1st of September and 31st of December of each season. An example of an annotated image is shown in Fig. 5A. From the colony boundaries we computed the surface area covered by the colony (in m²) by perspective correction and projection using the intrinsic and extrinsic camera parameters such as focal length, elevation, tilt, and roll (*30*). A projected top view is shown in Fig. 5B. The extracted areas of the Pointe Géologie colony between September 1 and December 31, 2014 are shown in Fig. 5H.

Colony densities (average number of individuals (adults plus chicks) per area) for Atka Bay and Pointe Géologie are calculated from individual counts (weekly resolution) divided by colony area for each of the images. For time points where we had area measurements but not a corresponding individual count, we linearly interpolated the value from the two nearest available counts.

We extended our dataset of colony area with satellite image based measurements of colony area from Coulman Island (12 days), Atka Bay (19 days), and Stancomb-Wills Glacier (18 days), between September 10 and December 11, 2011 published in (*22*).

### Meteorological data

The meteorological data for Pointe Géologie and Atka Bay (seasons 2018-2020) were recorded every minute by the meteorological observatory at Dumont d’Urville station (operated by Météo France) and the meteorological observatory at Neumayer station. The meteorological data for Coulman Island, Atka Bay (season 2011), and Stancomb-Wills Glacier stem from the ECMWF model “The ERA5 global reanalysis”, which provides 1h temporal and 1km² spatial resolution (*31*) and is available online (https://www.ecmwf.int/en/forecasts/dataset/ecmwf-reanalysis-v5).

### Phenological model

We developed a mechanistic phenological model to describe the temporary fluctuations of the number penguins at the colony over the course of a breeding season.

The most important model parameter is the number breeding males and females. This number is assumed to remain constant during a breeding cycle but can change between years. Additionally, the colony consists of a number of non-breeding animals. The model then describes the temporary fluctuations of the probability that male and female breeding animals or non-breeding animals are present at the colony. These probabilities are computed from the following phenological events:

- arrival of the animals (breeders and non-breeders) at the colony site at the beginning of the breeding season,
- departure of the females after courtship and laying an egg, and departure of the non-breeders after unsuccessful courtship,
- return of the females for chick feeding and departure of the males for foraging at sea after hatching of the chicks,
- subsequent (2x, according to (*18*)) parental switching between feeding and foraging, departure of both parents after their chick becomes thermally independent,
- subsequent (7x, according to (*18*)) feeding and foraging trips (females and males) until fledging of the chicks.

Note, that the number of foraging trips (2x during guarding phase and 7x during crèching phase) represent the average over the whole colony. We could not introduce the trip number as a free parameter of the model without losing numeric stability.

In the model, these events are described as probability densities with Gaussian shape that sets the mean time point and the standard deviation (temporal spreading) of the event (see Fig. 7A). Each event is specific for male, female, or non-breeding animals. The amplitude of the probability density for each event is chosen so that its integral corresponds to the total number of all animals of a specific sex (or of the total number of all non-breeding animals) that participate in the event. For example, as chicks die, the number of parents returning to the colony for feeding decreases, which we model by a smaller amplitude of the probability density for events describing the feeding trips.

**Fig. 7.**
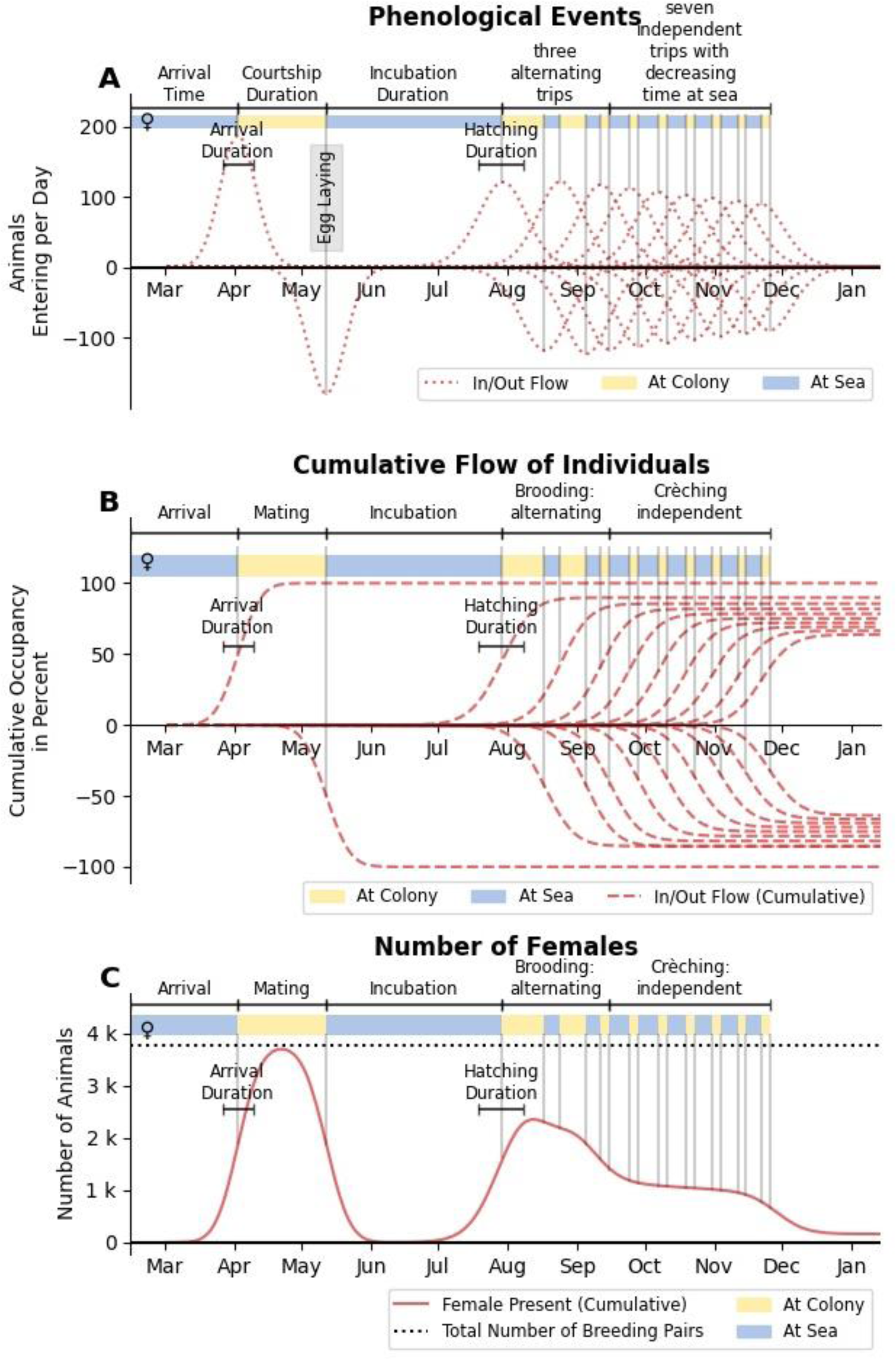
Illustration of the breeding cycle and the phenological model. (based on data from 2012 at Pointe Géologie). The model describes the number of individuals present at the colony as a result of phenological events that bring individuals to return or leave the colony site. This pattern of presence and absence is represented by colored bars (yellow: at the colony, blue: at sea) in each of the plots. The model assumes that the probability densities for individual animals of returning and leaving the colony are normally distributed. The mean and width of each of these distributions are model parameters. (**A**) Probability density distribution of all phenological events included in the model. Arrivals are indicated by positive values, departures by negative values. The width of the distributions indicates the distributions of individual arrival or departure times. The height corresponds to the number of individuals participating in the event. (**B**) Cumulative probabilities (computed by integrating the probability density distributions) indicate the number of individuals over time per event as a percentage of the total number of breeding pairs. Positive values indicate arrivals, negative values indicate departures. Note that the number of adults decreases over time due to loss of eggs or chicks. (**C**) Projected number of females present at the colony, computed as the sum of the cumulative probabilities in (B) multiplied by the total number of breeding pairs (dotted black line).

The number of chicks, their hatching and fledging time are not independently modeled. Rather, the hatching time coincides with the first return of the females (Stonehouse 1953; Prevost and Sapin-Jaloustre 1965). The fledging time coincides with the last parental departure. The number of hatching chicks is determined by the amplitude of the event describing the first return of the females, and the number of fledging chicks is determined by the amplitude of the last parental departure.

The number of male, female, or non-breeding animals is computed from the integration of the event probabilities over time (see Fig. 7B & 7C). The number of chicks is computed from the integration of the female event probabilities over time, starting with their first return after incubation.

The model parameters (time point, standard deviation, and amplitude of each event) are determined by a least squares fit of the model predictions to data. Since observers usually cannot distinguish between male and female animals, the sum of all adult animals is considered. For fitting the model, the number of chicks is not considered, however if this number is available from observations, it can be used for benchmarking the predictive power of the model.

The time points of the two parental switching (between feeding at the colony and foraging at sea) events for each sex, and the 7 consecutive feeding and foraging trips, are not free parameters. Rather, they are determined by the duration of a foraging trip at sea and the duration of the stay at the colony for feeding the chick. Furthermore, we assume that both the foraging and feeding durations decrease over time, in line with field observations (*16–21*), which we model by two linear functions (see Fig. 8). Furthermore, the duration of the first foraging trip of the females is assumed to be shorter than the first foraging trip of the males, also in line with field observations (*18, 33*).

**Fig. 8.**
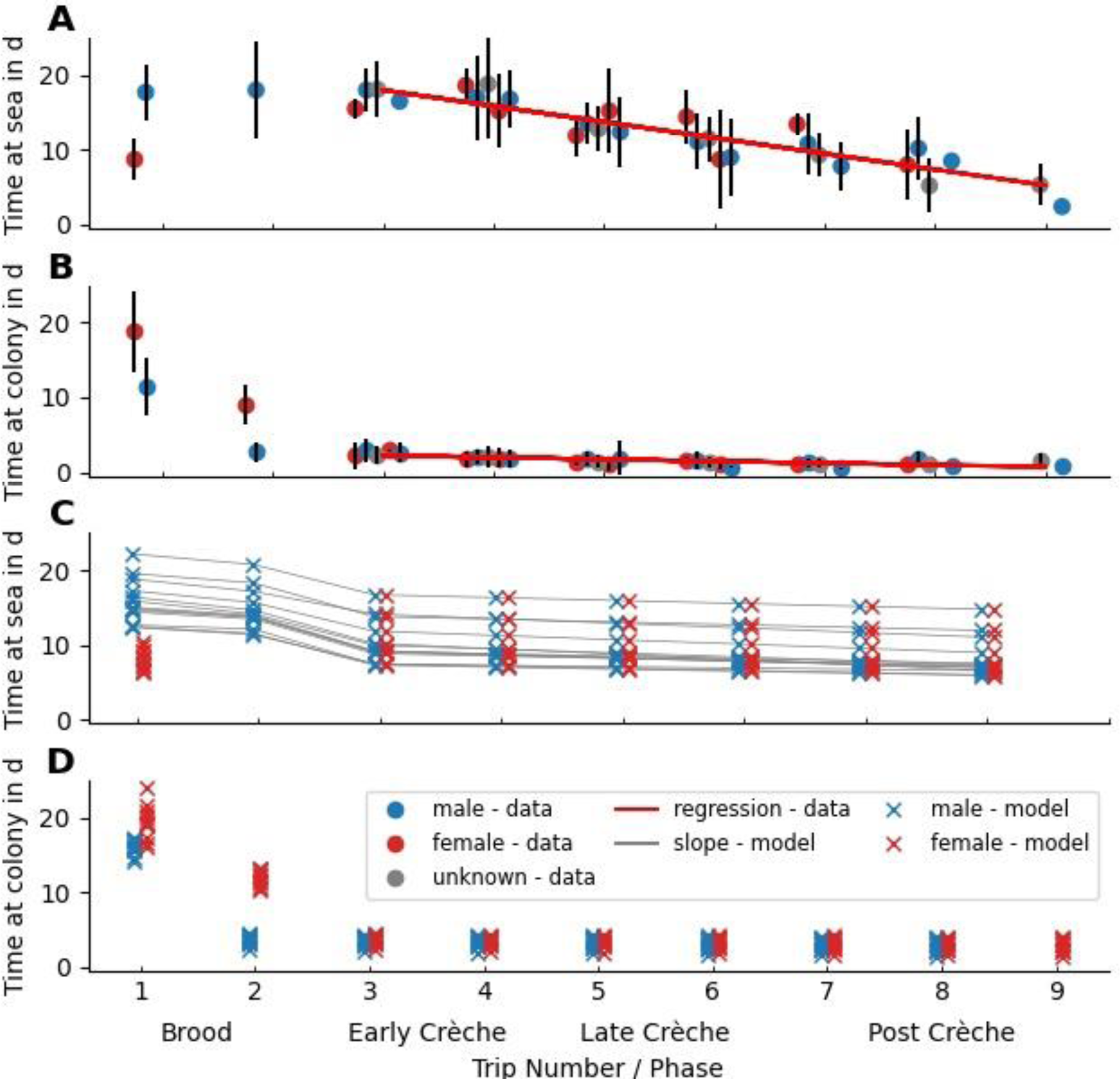
Individual trip durations during chick rearing phase. Time (in days, mean ± 1 standard deviation) a breeding emperor penguin spends at sea (**A**) or at the colony (**B**) at different stages of the breeding cycle (data from (*18*)). Red regression lines (**A**,**B**) show the decreasing trend in both time at sea and at the colony, after the initial brooding phase. (**C**) and (**D**) show the predicted time at sea and time at colony as predicted by the phenological model for 10 seasons (2012-2021) for Pointe Géologie and 3 seasons (2018-2020) for Atka Bay. Red, blue and gray dots denote the sex of the observed individuals (female/male/unknown). Gray lines indicate corresponding predictions for each season. Note that the model predicts large seasonal variations for the time at sea, but not for the time at colony.

Similarly, the standard deviations of all events are not free parameters but are constrained as follows: the standard deviation of arrival of all adult penguins, and the standard deviation of the first departure of females and non-breeders are the same, assuming that courtship is equally long for all breeders. The standard deviation of the first return of females and of all subsequent events are also equal and must be greater or equal than the standard deviation of arrival.

Taken together, the model has 14 free parameters (Table 1). We use Markov chain Monte Carlo (MCMC, see S1 in the Supplement) sampling to fit the model parameters to data (ground truth counts from manual observations). The best estimates of the model parameter for each colony and each season are listed in Table S3 in the Supplement.

### Windchill model

While they incubate their egg during the harsh Antarctic winter, emperor penguins conserve energy by forming tight groups, the so-called huddles (*24, 38*). The fraction of individuals of a colony that are currently in a huddle changes depending on an apparent (i.e. subjectively perceived) temperature. The apparent temperature depends on four environmental variables: ambient temperature (*T*), windspeed (*W*), solar radiation (*R*), and humidity (*H*). These variables linearly contribute to the apparent temperature T_a_, with linear factors *c_W_* (windchill factor), *c_R_* (solar radiation factor), *c_H_* (humidity factor), and a factor of unity for ambient temperature (Eq. 1) (*25*).

The model then describes the colony density as a sigmoidal function of the apparent temperature (Eq. 2). For very low apparent temperatures, the colony density tends to a maximum value of 12.8 animals / m^2^. This maximum density corresponds to a hexagonal packing of cylinders with a diameter of 30 cm. At very high apparent temperatures, the colony density tends to zero. The sigmoidal function has two free parameters, the critical temperature Tc at which the density is half of its maximum, and the steepness value b_0_ that describes how sensitively the animals respond to temperature changes.

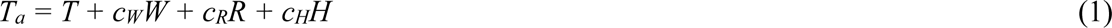

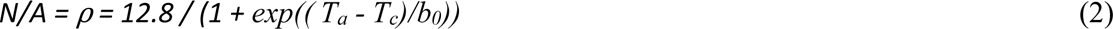

We fit the model parameters b_0_, c_W_, c_H_, c_R_, and T_c_ to the density data recorded at Pointe Géologie and Atka Bay between 2014 and 2020 for the months of September to December using Markov chain Monte Carlo sampling (see S1 in the Supplement). We can then predict the density of any colony at any given time point between September and December from weather data alone.

### Applying the phenological model to data from satellite images

When we fit the phenological model to count estimates from satellite images, problems may arise due to the small number of usable images for each season (typically less than 10 images, available only during the months from September to December). The fit of 14 free model parameters to such a small number of data points can lead to numerical instability and large confidence intervals. Therefore, we fix all except three model parameters to the average values that we obtained from the model fit to ground truth counts at Atka Bay and Pointe Géologie (see Table S3 in the Supplement). The time of arrival is chosen so that the date of female return aligns with the first sunrise after mid winter + 27.4 d. The only two free fit parameters are the number of breeding pairs and fledging success. This approach is justified by the fact that the phenology of the two colonies we have studied here (AB and PG) are similar except for the arrival time, as seen by the similar values of the model parameters and the absence of large systematic fluctuations during different years.

## Supporting information

Supplemental Information (General)

## Acknowledgements

This work was supported by the Deutsche Forschungsgemeinschaft (DFG) in the framework of the priority programme 1158 “Antarctic Research with comparative investigations in Arctic ice areas” by grants (FA336/5-1, ZI1525/3-1, ZI1527/7-1), by the Institut Polaire Français Paul-Emile Victor (IPEV) within the framework of the Project 137-ANTAVIA, by the Alfred-Wegener-Institut Helmholtz-Zentrum für Polar-und Meeresforschung (AWI) within the framework of the Projects SPOT and MARE, by the Centre Scientifique de Monaco with additional support from the LIA-647 and RTPI-NUTRESS (CSM/CNRS-UNISTRA), by the Centre National de la Recherche Scientifique (CNRS) through the Programme Zone Atelier Antarctique et Terres Australes (ZATA). We thank Météo France for the meteorological data of Dumont d’Urville. We are deeply grateful to all the wintering and summering members of projects IPEV 137, AWI-SPOT, AWI-MARE, and we also sincerely thank the IPEV and AWI logistics teams for their important and continued support in the field.

## Funding

German Research Foundation grant FA336/5-1 (BF)

German Research Foundation grant ZI1525/3-1 (DZ)

German Research Foundation grant ZI1527/7-1 (DZ)

National Science Foundation grant 2046437 (DZ)

Woods Hole Oceanographic Institution (DZ)

Institut Polaire Français Paul-Emile Victor (IPEV) 137-ANTAVIA (CLB)

Centre Scientifique de Monaco (CLB)

Centre National de la Recherche Scientifique (CLB)

CSM/CNRS-UNISTRA (LIA-647 and RTPI-NUTRESS) (CLB)

## Author Contributions

Conceptualization: AW, SR, BF, CLB, DZ

Methodology: AW, SR, BF, AM, CM, CLB, DZ

Data analysis: AW

Data collection: AW, SR, AH, TB, MB, CC, DC, RC, CE, BF, AK, AM, DM, JM, SP, ES, CLB, DZ

Supervision: BF, CLB, DZ

Writing—original draft: AW, SR, AH, BF, CLB, DZ

All authors contributed critically to the drafts and gave final approval for publication.

## Data and materials availability

All data are available in the main text or the supplementary materials.

## References

1. C. Barbraud, H. Weimerskirch, Emperor penguins and climate change. Nature. 411, 183–186 (2001).

2. J. Forcada, P. N. Trathan, Penguin responses to climate change in the Southern Ocean. Glob. Change Biol. (2009), doi:10.1111/j.1365-2486.2009.01909.x.

3. S. Jenouvrier, M. Holland, D. Iles, S. Labrousse, L. Landrum, J. Garnier, H. Caswell, H. Weimerskirch, M. LaRue, R. Ji, C. Barbraud, The Paris Agreement objectives will likely halt future declines of emperor penguins. Glob. Change Biol. 26, 1170–1184 (2019).

4. S. Jenouvrier, M. Holland, J. Stroeve, M. Serreze, C. Barbraud, H. Weimerskirch, H. Caswell, Projected continent-wide declines of the emperor penguin under climate change. *Nat*. Clim. Change. 4, 715–718 (2014).

5. P. T. Fretwell, P. N. Trathan, Discovery of new colonies by Sentinel2 reveals good and bad news for emperor penguins. *Remote Sens*. Ecol. Conserv. 7, 139–153 (2020).

6. P. N. Trathan, B. Wienecke, C. Barbraud, S. Jenouvrier, G. Kooyman, C. Le Bohec, D. G. Ainley, A. Ancel, D. P. Zitterbart, S. L. Chown, M. LaRue, R. Cristofari, J. Younger, G. Clucas, C.-A. Bost, J. A. Brown, H. J. Gillett, P. T. Fretwell, The emperor penguin - Vulnerable to projected rates of warming and sea ice loss. Biol. Conserv. 241, 108216 (2020).

7. D. Ainley, G. Ballard, L. K. Blight, S. Ackley, S. D. Emslie, A. Lescroël, S. Olmastroni, S. E. Townsend, C. T. Tynan, P. Wilson, E. Woehler, Impacts of cetaceans on the structure of Southern Ocean food webs. Mar. Mammal Sci. 26, 482–498 (2010).

8. A. D. Fraser, P. Wongpan, P. J. Langhorne, A. R. Klekociuk, K. Kusahara, D. Lannuzel, R. A. Massom, K. M. Meiners, K. M. Swadling, D. P. Atwater, G. M. Brett, M. Corkill, L. A. Dalman, S. Fiddes, A. Granata, L. Guglielmo, P. Heil, G. H. Leonard, A. R. Mahoney, A. McMinn, P. van der Merwe, C. K. Weldrick, B. Wienecke, Rev. Geophys., in press, doi:10.1029/2022RG000770.

9. Springing into summer | Arctic Sea Ice News and Analysis (2023), (available at https://nsidc.org/arcticseaicenews/2023/06/springing-into-summer/).

10. C. Le Bohec, J. D. Whittington, Y. Le Maho, “Polar Monitoring: Seabirds as Sentinels of Marine Ecosystems” in Adaptation and Evolution in Marine Environments, Volume 2: The Impacts of Global Change on Biodiversity, C. Verde, G. di Prisco, Eds. (Springer, Berlin, Heidelberg, 2013; 10.1007/978-3-642-27349-0_11), From Pole to Pole, pp. 205–230.

11. H. Weimerskirch, P. Jouventin, J. L. Mougin, J. C. Stahl, M. V. Beveren, Banding Recoveries and the Dispersal of Seabirds Breeding in French Austral and Antarctic Territories. Emu - Austral Ornithol. 85, 22–33 (1985).

12. S. Jenouvrier, M. Holland, J. Stroeve, C. Barbraud, H. Weimerskirch, M. Serreze, H. Caswell, Effects of climate change on an emperor penguin population: analysis of coupled demographic and climate models. Glob. Change Biol. 18, 2756–2770 (2012).

13. S. Richter, R. C. Gerum, W. Schneider, B. Fabry, C. Le Bohec, D. P. Zitterbart, A remote-controlled observatory for behavioural and ecological research: A case study on emperor penguins. Methods Ecol. Evol. (2018), doi:10.1111/2041-210X.12971.

14. B. Stonehouse, The emperor penguin (Aptenodytes forsteri, Gray): I. Breeding behaviour and development (1953).

15. J. Prevost, J. Sapin-Jaloustre, “Ecologie des Manchots Antarctiques” in Biogeography and Ecology in Antarctica, J. van Mieghem, P. van Oye, Eds. (Springer Netherlands, Dordrecht, 1965; 10.1007/978-94-015-7204-0_16), Monographiae Biologicae, pp. 551–648.

16. A. Ancel, G. L. Kooyman, P. J. Ponganis, J.-P. Gendner, J. Lignon, X. Mestre, N. Huin, P. H. Thorson, P. Robisson, Y. Le Maho, Foraging behaviour of emperor penguins as a resource detector in winter and summer. Nature. 360, 336–339 (1992).

17. G. L. Kooyman, T. G. Kooyman, Diving Behavior of Emperor Penguins Nurturing Chicks at Coulman Island, Antarctica. The Condor. 97, 536–549 (1995).

18. R. Kirkwood, G. Robertson, Seasonal change in the foraging ecology of emperor penguins on the Mawson Coast, Antarctica. Mar. Ecol. Prog. Ser. 156, 205–223 (1997).

19. I. Zimmer, R. P. Wilson, C. Gilbert, M. Beaulieu, A. Ancel, J. Plötz, Foraging movements of emperor penguins at Pointe Géologie, Antarctica. Polar Biol. 31, 229– 243 (2008).

20. S. Watanabe, K. Sato, P. J. Ponganis, Activity Time Budget during Foraging Trips of Emperor Penguins. PLoS ONE. 7, e50357 (2012).

21. A. Houstin, D. P. Zitterbart, A. Winterl, S. Richter, V. Planas-Bielsa, D. Chevallier, A. Ancel, J. Fournier, B. Fabry, C. L. Bohec, Biologging of emperor penguins— Attachment techniques and associated deployment performance. PLOS ONE. 17, e0265849 (2022).

22. S. Labrousse, D. Iles, L. Viollat, P. Fretwell, P. N. Trathan, D. P. Zitterbart, S. Jenouvrier, M. LaRue, Quantifying the causes and consequences of variation in satellite-derived population indices: a case study of emperor penguins. Remote Sens. Ecol. Conserv. 8, 151–165 (2021).

23. P. T. Fretwell, M. A. LaRue, P. Morin, G. L. Kooyman, B. Wienecke, N. Ratcliffe, A. J. Fox, A. H. Fleming, C. Porter, P. N. Trathan, An Emperor Penguin Population Estimate: The First Global, Synoptic Survey of a Species from Space. PLOS ONE. 7, e33751 (2012).

24. C. Gilbert, D. McCafferty, Y. Le Maho, J.-M. Martrette, S. Giroud, S. Blanc, A. Ancel, One for all and all for one: the energetic benefits of huddling in endotherms. Biol. Rev. Camb. Philos. Soc. 85, 545–69 (2010).

25. S. Richter, R. Gerum, A. Winterl, A. Houstin, M. Seifert, J. Peschel, B. Fabry, C. Le Bohec, D. P. Zitterbart, Phase transitions in huddling emperor penguins. J. Phys. Appl. Phys. 51 (2018), doi:10.1088/1361-6463/aabb8e.

26. R. Massom, K. Hill, C. Barbraud, N. Adams, A. Ancel, L. Emmerson, M. Pook, Fast ice distribution in Adélie Land, East Antarctica: interannual variability and implications for emperor penguins Aptenodytes forsteri. Mar. Ecol. Prog. Ser. 374, 243–257 (2009).

27. P. T. Fretwell, P. N. Trathan, Emperors on thin ice: three years of breeding failure at Halley Bay. Antarct. Sci. 31, 133–138 (2019).

28. A. Winterl, S. Richter, A. Houstin, A. P. Nesterova, F. Bonadonna, W. Schneider, B. Fabry, C. L. Bohec, D. P. Zitterbart, micrObs – A customizable time-lapse camera for ecological studies. HardwareX. 8 (2020), doi:10.1016/j.ohx.2020.e00134.

29. R. C. Gerum, S. Richter, B. Fabry, D. P. Zitterbart, *ClickPoints* : an expandable toolbox for scientific image annotation and analysis. Methods Ecol. Evol. 8, 750–756 (2017).

30. R. C. Gerum, S. Richter, A. Winterl, C. Mark, B. Fabry, C. Le Bohec, D. P. Zitterbart, CameraTransform: A Python package for perspective corrections and image mapping. SoftwareX. 10, 100333 (2019).

31. H. Hersbach, B. Bell, P. Berrisford, S. Hirahara, A. Horányi, J. Muñoz-Sabater, J. Nicolas, C. Peubey, R. Radu, D. Schepers, A. Simmons, C. Soci, S. Abdalla, X. Abellan, G. Balsamo, P. Bechtold, G. Biavati, J. Bidlot, M. Bonavita, G. Chiara, P. Dahlgren, D. Dee, M. Diamantakis, R. Dragani, J. Flemming, R. Forbes, M. Fuentes, A. Geer, L. Haimberger, S. Healy, R. J. Hogan, E. Hólm, M. Janisková, S. Keeley, P. Laloyaux, P. Lopez, C. Lupu, G. Radnoti, P. Rosnay, I. Rozum, F. Vamborg, S. Villaume, J. Thépaut, The ERA5 global reanalysis. Q. J. R. Meteorol. Soc. 146, 1999– 2049 (2020).

32. S. Labrousse, A. D. Fraser, M. Sumner, F. Le Manach, C. Sauser, I. Horstmann, E. Devane, K. Delord, S. Jenouvrier, C. Barbraud, Landfast ice: a major driver of reproductive success in a polar seabird. Biol. Lett. 17, 20210097 (2021).

33. T. D. Williams, The Penguins: Spheniscidae (Oxford University Press, 1995).

34. F. Ramírez, A. Tarroux, J. Hovinen, J. Navarro, I. Afán, M. G. Forero, S. Descamps, Sea ice phenology and primary productivity pulses shape breeding success in Arctic seabirds. Sci. Rep. 7, 4500 (2017).

35. J. Durant, G. Ottersen, N. C. Stenseth, Climate and the match or mismatch between predator requirements and resource availability. Clim. Res. 33, 271–283 (2007).

36. D. H. Cushing, “Plankton Production and Year-class Strength in Fish Populations: an Update of the Match/Mismatch Hypothesis” in Advances in Marine Biology, J. H. S. Blaxter, A. J. Southward, Eds. (Academic Press, 1990; https://www.sciencedirect.com/science/article/pii/S0065288108602023), vol. 26, pp. 249–293.

37. D. H. Cushing, The Regularity of the Spawning Season of Some Fishes. ICES J. Mar. Sci. 33, 81–92 (1969).

38. Y. Le Maho, The Emperor Penguin: A Strategy to Live and Breed in the Cold: Morphology, physiology, ecology, and behavior distinguish the polar emperor penguin from other penguin species, particularly from its close relative, the king penguin. Am. Sci. 65, 680–693 (1977).

39. J. Salvatier, T. V. Wiecki, C. Fonnesbeck, “Probabilistic programming in Python using PyMC3” (e1686v1, PeerJ Inc., 2016), doi:10.7287/peerj.preprints.1686v1.

40. M. D. Hoffman, A. Gelman, The No-U-Turn Sampler: Adaptively Setting Path Lengths in Hamiltonian Monte Carlo. J. Mach. Learn. Res. 15, 1351–1381 (2014).

41. A. Gelman, D. B. Rubin, Inference from Iterative Simulation Using Multiple Sequences. Stat. Sci. 7, 457–472 (1992).

