## Supplemental Information (General) for "Remote sensing of emperor penguin abundance and breeding success"

### Supplement

**S1 MCMC-Sampling**

**Phenological Model**

We inferred the values for the model parameters using Bayesian inference based on Markov Chain Monte Carlo (MCMC) sampling implemented in Python (*39*, *40*). The prior distributions for all parameters were uniform in between the boundaries given in Table 1. The counted number of individuals is subject to a statistical measurement error and accordingly follows a distribution. We assumed that the main counting error is relative: the human observers miscount by a ratio rather than a fixed number. Consequently and in accordance with literature (*22*), we chose the log-normal distribution as the individual counts error distribution, because its width scales approximately linear with the mean value. We tackled numerical issues at very low numbers (<100 individuals) by adding a constant offset number of non-breeding individuals that could be present regardless of the phenology. The width of the log-normal distribution and the offset number are free parameters. We chose an exponential distribution as the prior for both parameters to ensure minimization of error distribution width during the sampling process. Table 1 contains all free parameters of the model, the numerical range, prior distribution, and a brief description.

We used the 511 counts of adult individuals to optimize the parameters of our model. We sampled the 208 parameters (16 per season, 10 seasons at Point Géologie, 3 seasons at Atka Bay) of the model with the No-U-Turn-Sampler (*40*) for 200 tuning and 200 draw iterations in 8 independent chains. The parameters and therefore sampling statistics for different seasons and colonies are independent by model definition. The tuning samples were discarded after the sampling process. We use the pooled 1600 draw samples to compute means and errors for all parameters and other result plots. We computed a R-hat statistics (*41*) of less than 1.02 for each of the parameters. Consequently, the sampling has converged. S5 in the Supplement shows a trace plot containing the trace and kernel density estimate for all parameters.

##### Windchill Model

We used MCMC sampling and Bayesian statistics to infer the parameters of the density model while only using the measured areas and meteorological data as inputs. We applied non-informative normal distributions as priors for the linear factors and the transition temperature ($c_T = 1 / b_0$, $c_W$, $c_R$, $c_H$, $T_c$). Again, we modeled the error distribution of the density ($\rho$) as a log-normal distribution with standard deviation $\sigma$. Analogous to the phenological model, we choose an exponential distribution as the prior for $\sigma$.

#### S2 Kernel density estimate of phenological model parameters

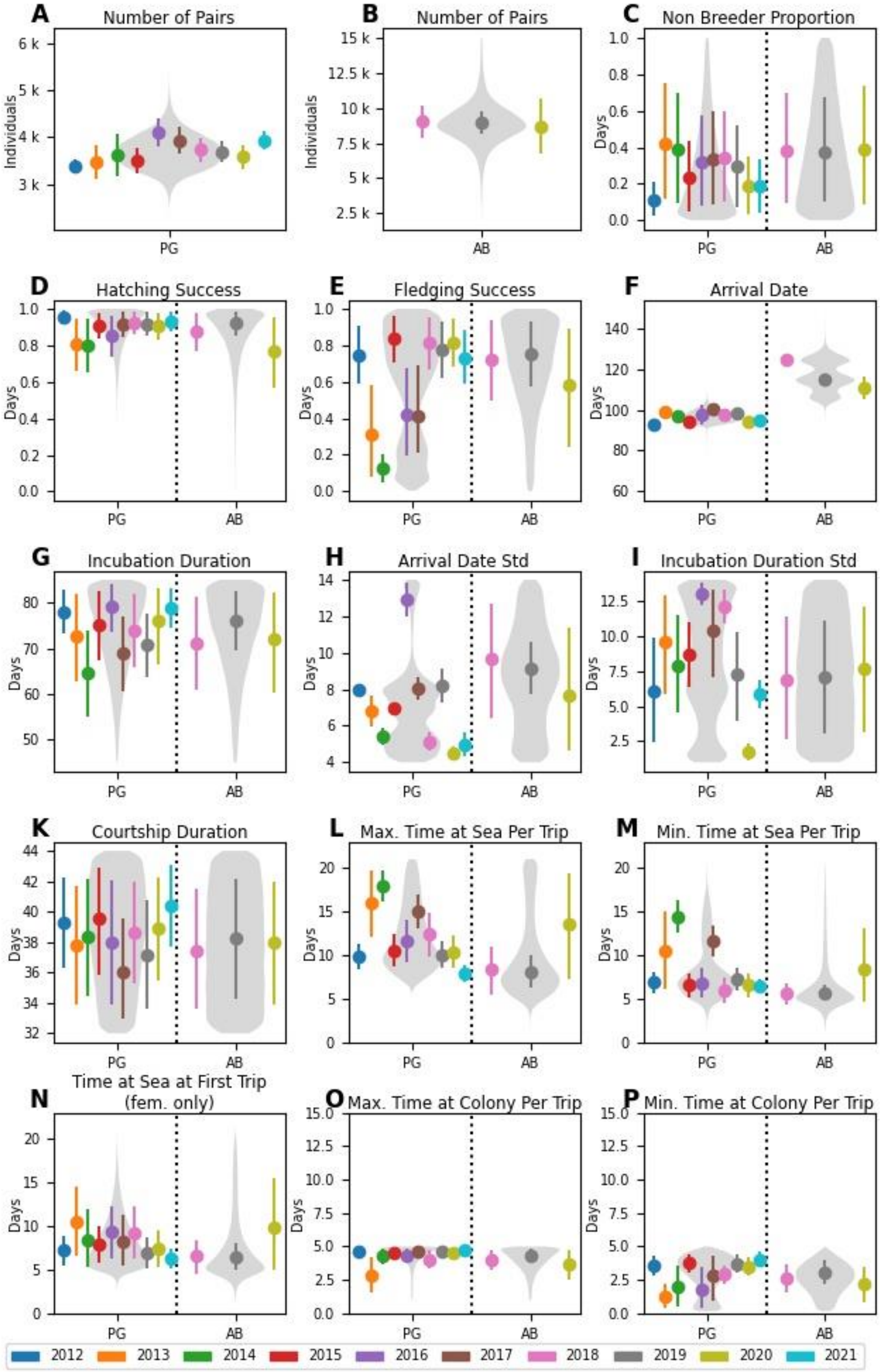


**Fig. S2. Distribution plots of the model parameters for Pointe Géologie and Atka Bay**

**colonies.** In each panel, the left part shows the parameters for Pointe Géologie colony (PG) and the right part for Atka Bay colony (AB). The points show the parameter value for each year, while the violin plots show the kernel density estimation for the parameter.

#### S3 Numerical Values of Phenological Parameters

| Colony | Season | Number of Pairs | Non Breeder Proportion | Hatching Success | Fledging Success | Arrival Date | Arrival Date Std | Courtship Duration | Female Absence Duration | Female Absence Duration Std | Max. Time at Colony Per Trip | Min. Time at Colony Per Trip | Max. Time at Sea Per Trip | Time at Sea at First Trip (fem. only) | Min. Time at Sea Per Trip | Relative Error | Offset Error |
|---|---|---|---|---|---|---|---|---|---|---|---|---|---|---|---|---|---|
| **Unit** | | 1 | 1 | 1 | 1 | d | d | d | d | d | d | d | d | d | d | 1 | 1 |
| **Pointe Géologie** | **2012** | 3380. | 0.10 | 0.95 | 0.70 | 92.80 | 8.00 | 39.00 | 78.00 | 6.00 | 4.60 | 3.60 | 9.80 | 7.20 | 6.90 | 0.18 | 9.70 |
| | **2013** | 3500. | 0.40 | 0.80 | 0.30 | 99.30 | 6.80 | 38.00 | 73.00 | 10.00 | 2.80 | 1.30 | 16.00 | 11.00 | 10.00 | 0.31 | 12.00 |
| | **2014** | 3600. | 0.40 | 0.80 | 0.12 | 96.70 | 5.40 | 38.00 | 65.00 | 8.00 | 4.30 | 2.00 | 17.90 | 8.00 | 14.40 | 0.50 | 1.40 |
| | **2015** | 3500. | 0.20 | 0.91 | 0.80 | 94.30 | 6.90 | 40.00 | 75.00 | 9.00 | 4.50 | 3.70 | 10.50 | 8.00 | 6.60 | 0.33 | 1.20 |
| | **2016** | 4100. | 0.30 | 0.80 | 0.40 | 98.00 | 12.90 | 38.00 | 79.00 | 13.00 | 4.40 | 1.80 | 12.00 | 9.00 | 6.80 | 0.26 | 6.00 |
| | **2017** | 3900. | 0.30 | 0.92 | 0.40 | 100.60 | 8.00 | 36.00 | 69.00 | 10.00 | 4.60 | 2.80 | 14.90 | 8.00 | 11.60 | 0.29 | 1.80 |
| | **2018** | 3700. | 0.30 | 0.93 | 0.80 | 97.40 | 5.10 | 39.00 | 74.00 | 12.10 | 4.00 | 2.90 | 12.00 | 9.00 | 5.90 | 0.28 | 1.60 |
| | **2019** | 3700. | 0.30 | 0.92 | 0.80 | 98.00 | 8.20 | 37.00 | 71.00 | 7.00 | 4.60 | 3.70 | 10.10 | 6.90 | 7.30 | 0.23 | 33.00 |
| | **2020** | 3600. | 0.20 | 0.91 | 0.80 | 94.30 | 4.50 | 39.00 | 76.00 | 1.70 | 4.50 | 3.50 | 10.40 | 7.00 | 6.50 | 0.29 | 2.00 |
| | **2021** | 3900. | 0.20 | 0.93 | 0.70 | 95.10 | 5.00 | 40.00 | 79.00 | 5.90 | 4.70 | 4.00 | 7.90 | 6.30 | 6.40 | 0.21 | 1.90 |
| **Atka Bay** | **2018** | 9000. | 0.40 | 0.90 | 0.70 | 125.00 | 10.00 | 37.00 | 71.00 | 7.00 | 4.00 | 2.60 | 8.00 | 7.00 | 6.00 | 0.33 | 1600.0 |
| | **2019** | 9000. | 0.40 | 0.92 | 0.70 | 115.00 | 9.10 | 38.00 | 76.00 | 7.00 | 4.30 | 3.10 | 8.00 | 6.50 | 5.70 | 0.34 | 190.00 |
| | **2020** | 9000 | 0.4 | 0.8 | 0.6 | 110.0 | 8.0 | 38.0 | 72.0 | 8.0 | 3.7 | 2.2 | 14.0 | 10.0 | 8.0 | 0.6 | 700.0 |

#### S4 Individual Fits

Attach pdf files.

### S5 Traceplots

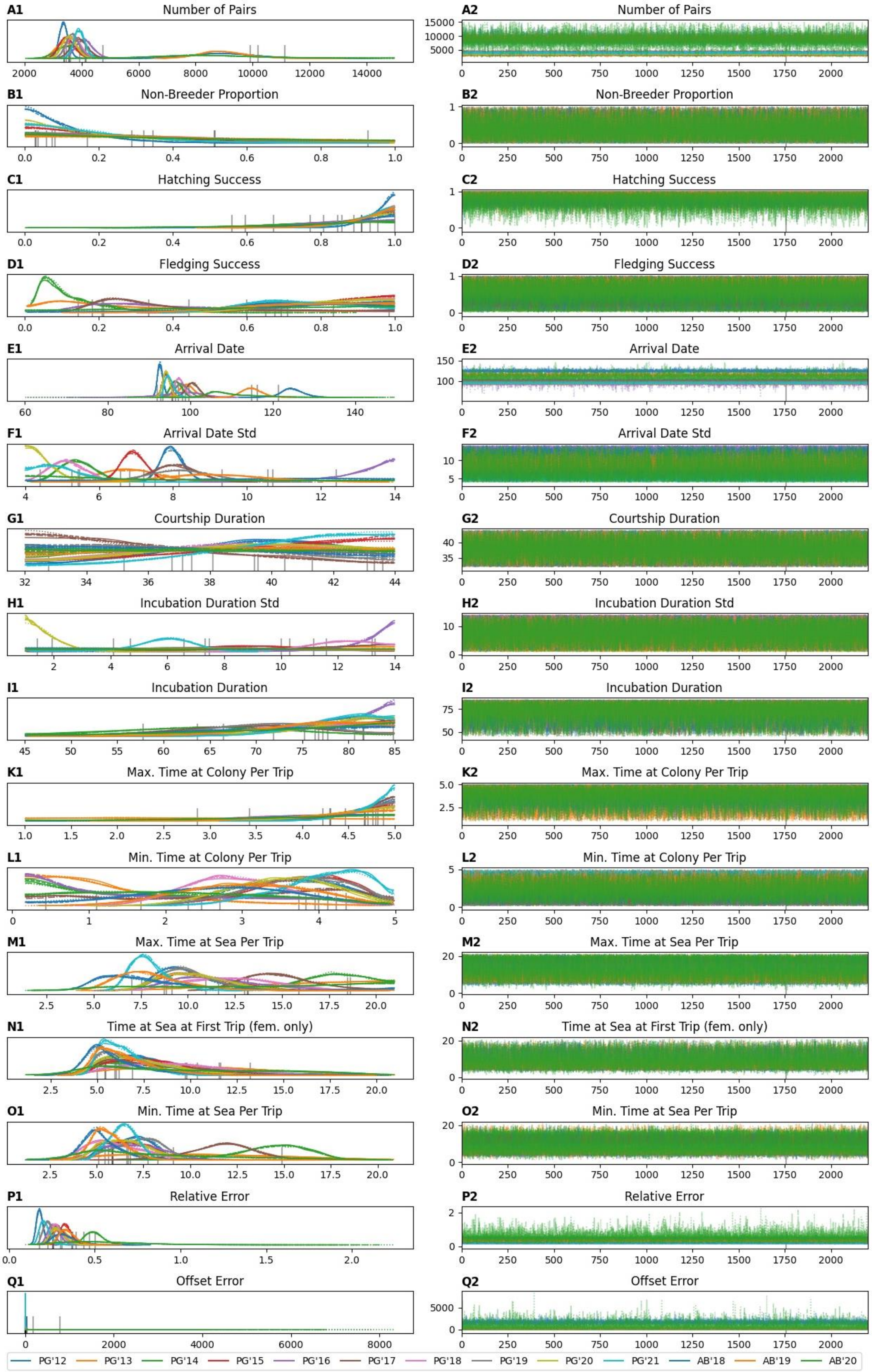

**Fig. S5.: Plot of kernel density estimates and traces from the sampling process.** Every row shows one parameter of the sampling process. The left column shows the kernel density estimates: On the X-axis the parameter value on the Y-axis the probability of that value. The right column shows the trace of the sampling process (after the removing the tuning samples): The X-axis shows the sample index from 0-2400. TheY-axis shows the parameter value associated with that sample. Different colors indicate different seasons. Different line styles indicate different chains (1 to 4) in the sampling process.

### S6 Correlation of Model Parameters and Breeding Success

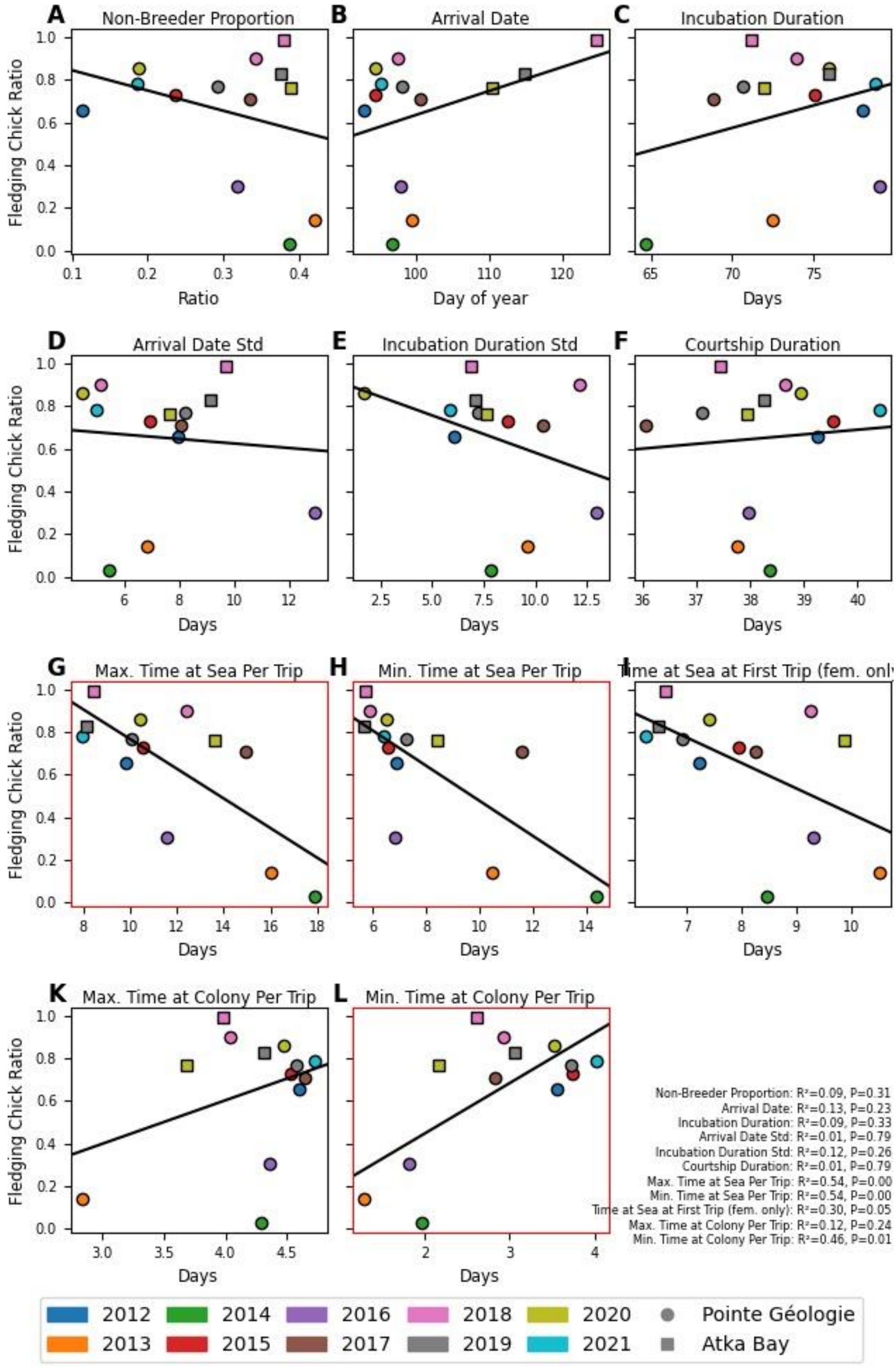

**Fig. S6.: Correlation of model parameters with breeding success.** The panels show the average predicted parameter value on X-axis and the ratio of fledging chicks to the number of breeders from manual observations on the Y-axis. Points show PG, squares show Atka Bay data. Colors indicate the year. The black line shows a linear regression fit performed on the data in the plot. We find significant correlation for the parameters $c_{min}$, $s_{max}$, and $s_{min}$ indicated by a red outline of the plot box.

##### S7 Satellite Data Based Fits

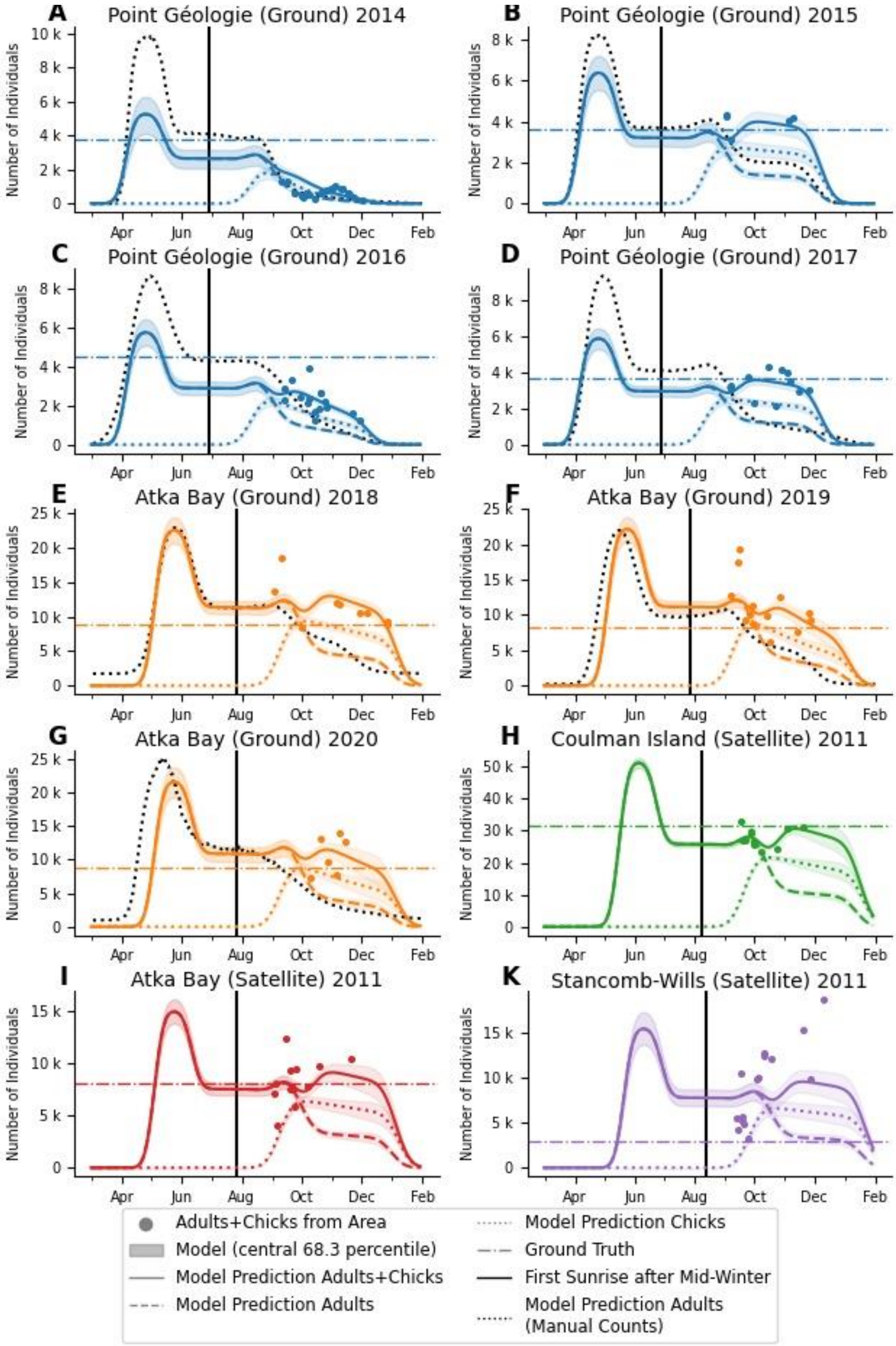


**Fig. S7.: Model predictions for satellite based data.** The plots show the total number of individuals per breeding season and colony. The points show the total number of individuals calculated by multiplying the colony covered area with the density predicted from meteorological values with the windchill model. Colony covered areas are measured either from ground (PG 2014-2017, AB 2018-2020) or from satellite (CI 2011, AB 2011, SW 2011) images. The colored dashed, dotted and solid lines represent the average number of adults, chicks and animals in total as predicted by the model based on the measured values represented in dots. The shaded areas indicate the 1-sigma interval (central 68.3 percentile). The dotted black line shows the number of adults predicted from a model fitted to manual count data including the whole season from March to January the following year (as in Fig. 2 C-F and Fig. 2 K-M). The horizontal dash-dotted lines show ground truth values for the total number of breeding pairs from manual observations (PG, AB 2018-2020) or from (*23*) (CI 2011, AB 2011, SW 2011). The vertical solid line shows the first sunrise after mid winter at the respective colony locations. According to our findings for PG and AB we used this date to align the phenological patterns of the other colonies. Colors indicate the colony and data source (ground or satellite based)

#### S8 Calculation of statistical metrics

##### R2-score

We calculate the R2 score as follows from two arrays x, and y containing the true and respective predicted values:

$u = \sum_i^N (x - y)^2$

$\mu = \sum_i^N x/N$

$v = \sum_i^N (x - \mu)^2$

R2 = 1 - u / v

##### Statistical significance of inter-colony variability

We calculate the average parameter value for each season/colony combination. Then we pool the Atka Bay (AB) and Pointe Géologie (PG) values and perform a two-sided Mann-Whitney U statistics (using the python package scipy version 1.7.3). If the resulting p-value is smaller than 0.05, we accept the parameter as colony-variant.

##### Statistical significance of inter-year variability

For each parameter set from the sampling chain, we calculate the pairwise differences of annual parameter values. This yields 45 distributions of parameter differences for the 10 years from Pointe Géologie and 3 distributions for the 3 years from Atka Bay. In these distributions zero equals no difference between years and a symmetric distribution around zero indicates no inter year variance. We count the number of samples left of zero and right of zero. We use Bonferroni-corrected significance values of 0.05/45 for Pointe Géologie and 0.05/3 for Atka Bay. If less than a significant portion of samples are on one side of zero, we accept the pair to be significantly different. If one of the inter year pairs for one colony is significantly different, we call the parameter inter-year variable at the respective colony.

We perform this test independently for Atka Bay and Pointe Géologie parameter sets.

**S9 Phenological Model Explicit Formulas**

We start the calculation of the number of individuals with a set of parameters as described above: BP, H, F, NB, $t_0$, $\Delta t_0$, m, b, $\Delta b$, $c_{max}$, $c_{min}$, $s_{max}$, $s_{fem}$, $s_{min}$

We create two arrays with 11 entries that contain the duration of foraging ($s_\mu$) and feeding ($c_\mu$) trips.

$s_\mu$ = (

11/11 $s_{max} + s_{min}$,

$s_{fem}$,

$s_{fem}$,

8/11 $s_{max} + s_{min} - s_{fem}$,,

7/11 $s_{max} + s_{min}$,

6/11 $s_{max} + s_{min}$,

5/11 $s_{max} + s_{min}$,

4/11 $s_{max} + s_{min}$,

3/11 $s_{max} + s_{min}$,

2/11 $s_{max} + s_{min}$,

1/11 $s_{max} + s_{min}$,

0/11 $s_{max} + s_{min}$,

)

$c_\mu$ = (

9/9 $c_{max} + c_{min}$,

9/9 $c_{max} + c_{min}$,

8/9 $c_{max} + c_{min}$,

7/9 $c_{max} + c_{min}$,

6/9 $c_{max} + c_{min}$,

5/9 $c_{max} + c_{min}$,

4/9 $c_{max} + c_{min}$,

3/9 $c_{max} + c_{min}$,

2/9 $c_{max} + c_{min}$,

1/9 $c_{max} + c_{min}$,

0/9 $c_{max} + c_{min}$,

)

We create two arrays with 26 entries that contain the mean ($t_{\mu,diff}$) and standard deviation ($t_{\sigma,diff}$) of the durations between every individual in or out event:

$t_{\mu,diff} = (t_0, m, b, s_{\mu,0}, c_{\mu,0}, s_{\mu,1}, c_{\mu,1}, \ldots s_{\mu,10}, c_{\mu,10}, c_{min})$

$t_{\sigma,diff} = (\Delta t_0, 0, \Delta b, 0, \ldots ,0)$

We arrive at the absolute mean and standard deviation of the event timings by applying a cumulative sum:

$t_\mu = \text{cumsum}(t_{\mu,diff})$

$t_\sigma = \text{cumsum}((t_{\sigma,diff})^2)^{0.5}$

With the sampling time T (in our case every day of the breeding season between March 1 and February 28 the next year.) we calculate the proportion $P_i(T)$ of individuals that have undergone a certain in/out event $t_{\mu,i}$ (i ranging between 0 and 25):
$P_i(T)$ = Errf( $(T-t_{\mu,i})/\ t_{\sigma,i}$ ) with being Errf(x) the normalized error function at x

We create 4 arrays ($f_M$, $f_F$, $f_{NB}$, $f_C$) with 26 entries containing the proportion of individuals that will take part in each of those events. This proportion depends on the type of individual (male breeder, female breeder, non breeder, visible chick) and the fraction parameters NB, H, and F. Negative numbers denote outgoing individuals.

$f_M$ = (1, 0, -1, 0, H, 0, 0, -H, 0, 0, $F^{1/7}$, $-F^{1/7}$, $F^{2/7}$, $-F^{2/7}$, $F^{3/7}$, $-F^{3/7}$, $F^{4/7}$, $-F^{4/7}$, $F^{5/7}$, $-F^{5/7}$, $F^{6/7}$, $-F^{6/7}$, $F^{7/7}$, $-F^{7/7}$, 0, 0)
$f_F$ = (1, -1, 1, H-1, O, -H, H, 0 -H, 0, $F^{1/7}$, $-F^{1/7}$, $F^{2/7}$, $-F^{2/7}$, $F^{3/7}$, $-F^{3/7}$, $F^{4/7}$, $-F^{4/7}$, $F^{5/7}$, $-F^{5/7}$, $F^{6/7}$, $-F^{6/7}$, $F^{7/7}$, $-F^{7/7}$, 0, 0)
$f_{NB}$ = (1, -1, 0, …, 0)
$f_C$ = (0, 0, 0, 0, 0, 0, 0, 0, 0, 0, $H-F^{1/7}$, $F^{2/7}-F^{1/7}$, 0, $F^{3/7}-F^{2/7}$, 0, $F^{4/7}-F^{3/7}$, 0, $F^{5/7}-F^{4/7}$, 0, $F^{6/7}-F^{5/7}$, 0, $F^{7/7}-F^{6/7}$, 0, 0, 0, -F)

Now we calculate the number of males ($N_M$), females ($N_F$), visible chicks ($N_C$) and non-breeders ($N_{NB}$) at sampling time T by multiplying $f_{X,i}$ and $P_i$ and summing over all events i (X is one of M,F,NB,C).
$N_M(T) = \sum_i f_{M,i}\, P_i(T)$
$N_F(T) = \sum_i f_{F,i}\, P_i(T)$
$N_{NB}(T) = \sum_i f_{NB,i}\, P_i(T)$
$N_C(T) = \sum_i f_{C,i}\, P_i(T)$

Finally, we calculate the number of adults BP by adding $N_M$ and $N_F$

#### S10 Individual Counts at Atka Bay and Point Geologie

Attach Excel Files

#### S11 Breeding Phenology Event Observations and Predictions

N.A. = "Not Available"

| Season | | 2012 | 2013 | 2014 | 2015 | 2016 | 2017 | 2018 | 2019 | 2020 | 2021 |
|---|---|---|---|---|---|---|---|---|---|---|---|
| **Arrival** | **true** | 2012-03-26 | 2013-04-06 | 2014-03-31 | 2015-03-30 | 2016-03-31 | 2017-04-01 | 2018-04-12 | 2019-04-05 | 2020-03-31 | 2021-03-29 |
| | **predicted** | 2012-04-02 | 2013-04-10 | 2014-04-07 | 2015-04-05 | 2016-04-07 | 2017-04-11 | 2018-04-08 | 2019-04-08 | 2020-04-04 | 2021-04-06 |
| **Female First Departure** | **true** | 2012-05-02 | 2013-04-30 | 2014-05-05 | 2015-05-12 | 2016-04-29 | 2017-04-26 | 2018-05-05 | 2019-05-03 | 2020-05-05 | 2021-05-02 |
| | **predicted** | 2012-05-12 | 2013-05-18 | 2014-05-16 | 2015-05-14 | 2016-05-15 | 2017-05-17 | 2018-05-17 | 2019-05-16 | 2020-05-13 | 2021-05-16 |
| **Female Return** | **true** | 2012-07-05 | 2013-07-26 | 2014-07-31 | 2015-07-18 | 2016-07-19 | 2017-07-27 | 2018-07-29 | 2019-07-26 | 2020-07-19 | 2021-07-26 |
| | **predicted** | 2012-07-29 | 2013-07-29 | 2014-07-19 | 2015-07-28 | 2016-08-02 | 2017-07-25 | 2018-07-30 | 2019-07-25 | 2020-07-28 | 2021-08-03 |
| **Male Return** | **true** | NA | 2013-08-19 | 2014-08-11 | 2015-08-04 | 2016-08-01 | 2017-08-06 | 2018-08-08 | 2019-08-08 | 2020-08-06 | 2021-08-09 |
| | **predicted** | 2012-08-12 | 2013-08-17 | 2014-08-10 | 2015-08-12 | 2016-08-18 | 2017-08-14 | 2018-08-15 | 2019-08-09 | 2020-08-11 | 2021-08-15 |
| **Emancipation** | **true** | 2012-08-23 | 2013-08-19 | 2014-08-16 | 2015-08-20 | 2016-08-17 | 2017-08-19 | 2018-08-17 | 2019-08-15 | 2020-08-09 | 2021-08-16 |
| | **predicted** | 2012-08-28 | 2013-09-02 | 2014-08-27 | 2015-08-29 | 2016-09-05 | 2017-08-31 | 2018-09-01 | 2019-08-25 | 2020-08-28 | 2021-08-31 |
| **Fledging** | **true** | 2012-12-14 | NA | 2015-01-07 | NA | 2016-12-30 | 2017-12-27 | 2018-12-04 | NA | 2020-12-03 | 2021-12-02 |
| | **predicted** | 2012-12-06 | 2013-12-28 | 2015-01-25 | 2015-12-09 | 2016-12-08 | 2018-01-12 | 2018-12-07 | 2019-12-06 | 2020-12-05 | 2021-12-04 |

#### S12 Numerical Values of Windchill Model

| | cT | cW | cR | cH | Tc | sigma |
|---|---|---|---|---|---|---|
| **mean** | -0.023845 | 0.079986 | -0.000767 | 0.008434 | -4.903825 | 0.731349 |
| **std** | 0.003476 | 0.007152 | 0.000133 | 0.001784 | 0.909726 | 0.023509 |

#### S13 Numerical Values of Phenological Parameters of Satellite Model

| **colony** | **unit** | **Point Géologie (Ground)** | | | | **Atka Bay (Ground)** | | | **Coulman Island (SAT)** | **Atka Bay (SAT)** | **Stancomb-Wills (SAT)** |
|---|---|---|---|---|---|---|---|---|---|---|---|
| **Season** | | **2014** | **2015** | **2016** | **2017** | **2018** | **2019** | **2020** | **2011** | **2011** | **2011** |
| **Number of Pairs** | 1 | 2800 | 3200 | 2900 | 3000 | 11200 | 11000 | 10800 | 25600 | 7300 | 7700 |
| **Number of Pairs (Year)** (23) | 1 | - | - | - | - | - | - | - | 25298 (2009) | 9657 (2009) | 5455 (2009) |
| **Fledging Success** | 1 | 0.09 | 0.80 | 0.37 | 0.80 | 0.70 | 0.50 | 0.60 | 0.80 | 0.80 | 0.80 |
| **Relative Error** | 1 | 0.62 | 0.23 | 0.25 | 0.25 | 0.19 | 0.27 | 0.27 | 0.09 | 0.27 | 0.46 |
| **Offset Error** | 1 | 10.00 | 11.00 | 10.00 | 10.00 | 10.00 | 10.00 | 10.00 | 10.00 | 10.00 | 10.00 |

#### S14 Definition and Calculation of "First Sunrise after mid winter"

We used the python package astropy (https://www.astropy.org/, version 5.2.2) to calculate the sun's elevation at noon throughout the year at the respective colony locations. We define the first sunrise after mid winter as the first day with a noon elevation over 0° after the winter solstice.

#### S15 Conversion Factor

In satellite based surveys, colony covered area is not converted to a total number of individuals on the day of image recording, but to a total number of breeders. For this, the studies employ a combined conversion factor that corrects for phenological and windchill effects. The conversion factor is close to 1 (e.g 0.93 in (*23*)) We compute a conversion factor *CF* from the number of breeding pairs from manual observations and the annual average of colony-covered area from ground based images as *CF = BP/A.* For the colonies and seasons in our study we find the following values:

CF = 5.98 (Pointe Géologie 2014), 0.90 (PG 2015), 2.13 (PG 2016), 1.14 (PG 2017), 0.78 (Atka Bay 2018), 0.80 (AB 2019), 0.82 (AB 2020).

#### S16 Preparation of Satellite Data

We use the windchill model to predict the density for each time point, where we measured the colony area. We calculate the number of individuals by multiplying the area with the density. In the case of ground-based images, we manually select the colony area by drawing a polygon. For spread-out colonies, average densities below 1 animal / m2 are possible. In the case of satellite images, by contrast, the colony area is extracted based on a pixel-wise classification, with each pixel being classified as “penguin” or “not-penguin”. As the pixels are approximately 1 m² in size, the minimum animal density associated with a pixel is therefore one animal/$m^2$. To account for this, we constrain the lower bound of the predicted density to 1 animal/m2.

#### S17 Parameter Distribution of Windchill Model

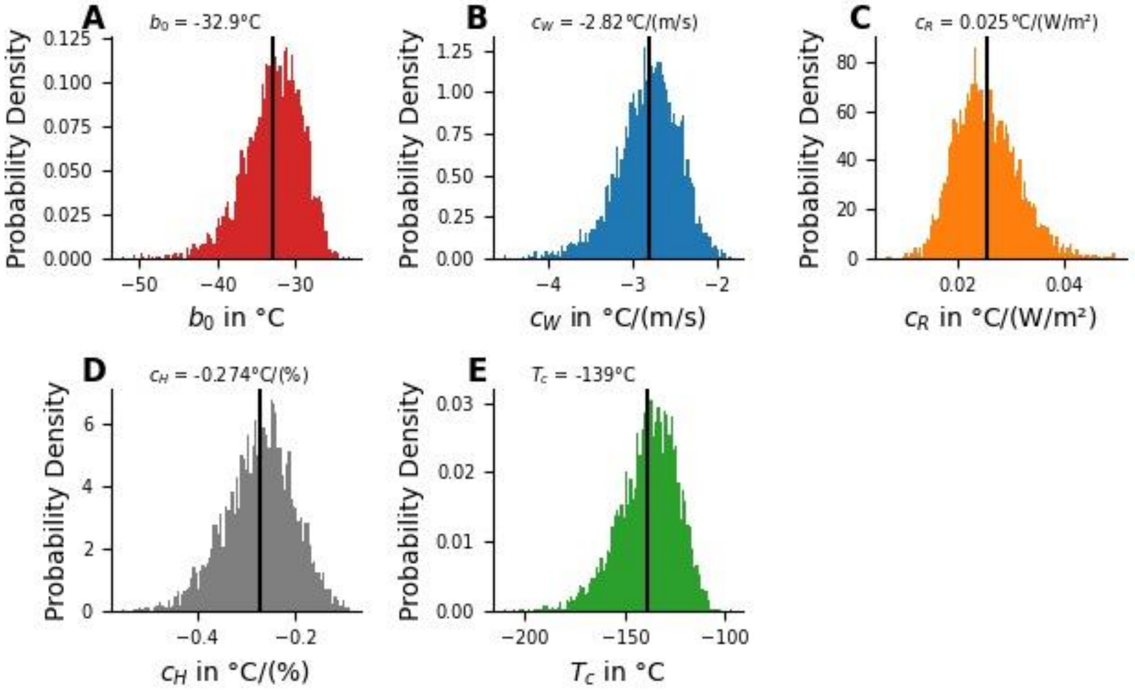


**Fig. S17.: Parameter distribution of windchill model.** The plots show the probability density for the 5 parameters of the windchill model $b_0$ (**A**), $c_W$ (**B**), $c_R$ (**C**), $c_H$ (**D**), and $T_C$ (**E**) as sampled by the windchill model. The solid vertical line and text in the figure show the average value for each parameter.

**S18 First sunrise after mid winter**

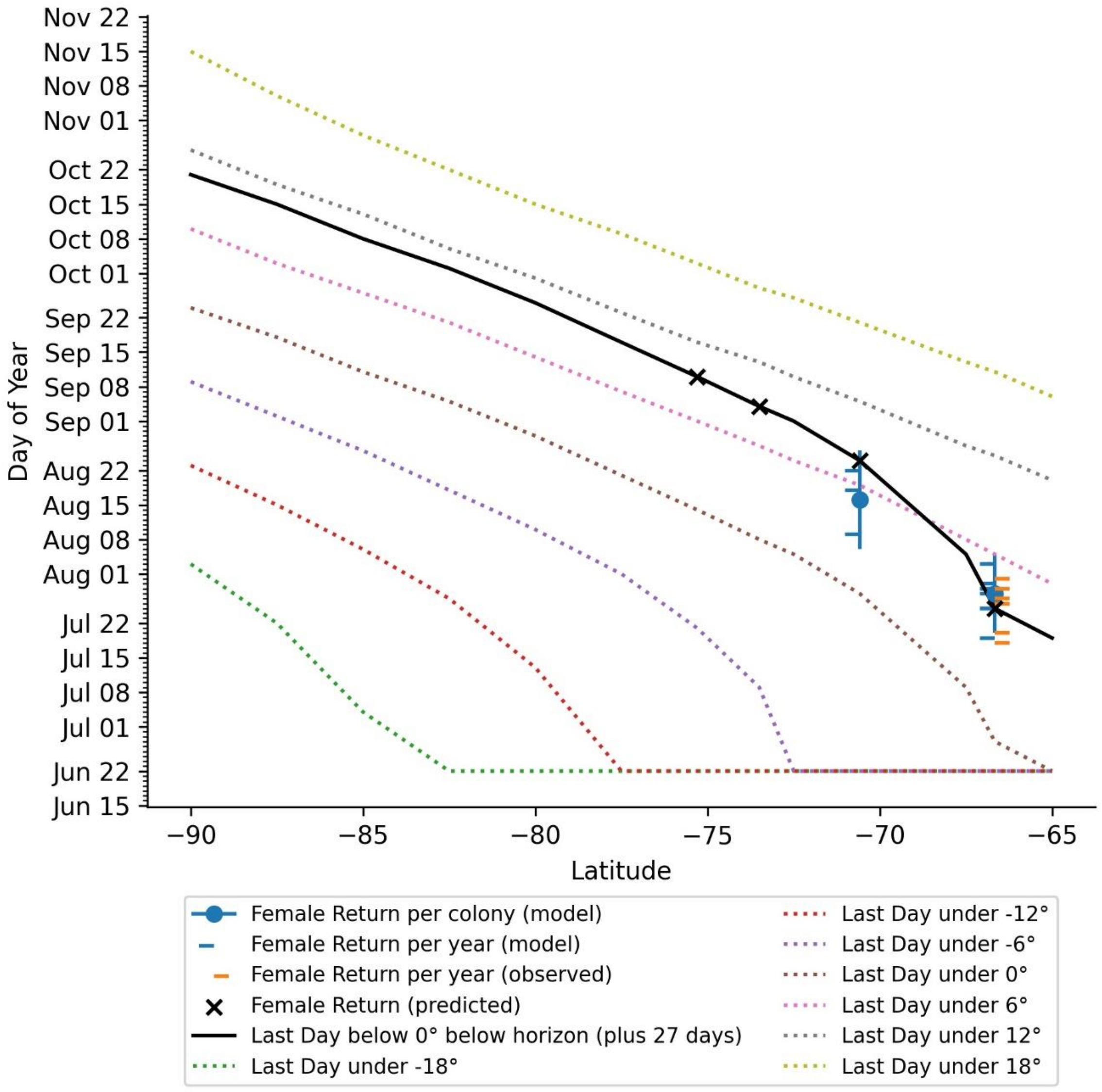


**Fig. S18.: Day of first sunrise after midwinter for different latitudes.** The plot shows latitude against the day of year. The blue dots and error bars show the average +/- standard deviation predicted day of female return as predicted by our phenology model. The blue left ticks show the predicted day of female return per season as predicted by the model. The orange right ticks show the day of female return as of manual observations. The dashed lines show the last day with a maximum solar elevation of -18°, -12°, -6°, 0°, 6°, 12°, and 18° at the respective latitude on the x-axis. Note: The lines for the low elevations (-18-0°) flatten out on June 21 for higher latitudes, because there are no days with such a small maximum sun elevation at those latitudes. The black line shows the day 27.4 days later than the last day with a maximum sun elevation smaller than 0°, which is our best fit for the prediction of the Female Return. The black crosses show the female return predicted for the latitudes of the 4 colonies in our study (Coulman Island, Stancomb-Wills, Atka Bay, and Pointe Géologie) according to this prediction.
